# Towards universal modeling of transcript isoform expression levels

**DOI:** 10.1101/2025.07.21.665977

**Authors:** Savio Ho-Chit Chow, Christina Huan Shi, Aniruddha Deshpande, Qin Cao, Kevin Y. Yip

## Abstract

A holy grail in computational biology is accurate modeling of transcript expression levels using epigenetic features, which would provide a quantitative way to study gene regulation in normal and disease states. Previous studies relied heavily on immortalized cell lines that exhibit properties different from cells in natural tissue environments. Most studies also quantified the expression of each gene by a single expression level, which fails to capture separate expression levels of different transcript isoforms of the same gene. In this study, making use of the latest large-scale dataset of paired transcriptomic and epigenomic data of human samples produced by the International Human Epigenome Consortium (IHEC), we computationally modeled the expression levels of individual transcript isoforms in 324 samples from 29 tissue types. We constructed the models using graph-based methods that integrate both location-specific epigenomic features and multiple types of gene-gene relationships. We found that to infer transcript isoform expression levels in a sample, a model that integrates information from many samples of other tissue types consistently outperforms a model trained on data from this sample itself, providing strong support that it is possible to construct a “universal” model that can accurately infer transcript isoform expression levels across tissue types.

## Introduction

Despite having almost identical genomic sequences, different cells in an organism can have very different gene expression profiles. A fundamental question in computational biology is that given various types of cell state measurements as input features, whether the expression level of each RNA product can be inferred by a fixed “universal” computational model that works for all tissue types, cell types, and cell states.

Previous studies have attempted to answer this question using different types of cell state measurements and modeling methods. In terms of cell state measurements, most of these studies considered transcription factor binding ^1–3^, histone modifications ^3–9^, DNA methylation ^10–12^, and chromatin accessibility ^3,5,13,14^. To turn each type of data into features to infer the expression level of a gene, aggregated signals were typically calculated at regions proximal to the gene such as its promoter ^2,4–12,15^ and gene body regions ^2,5,10,11^. Some studies have also considered these signals at more distal cis-regulatory elements (CREs), such as enhancers ^3,9,14,16,17^. There were also studies that took DNA sequences as input features, alone ^18,19^ or together with epigenomic features ^3^.

In terms of modeling methods, early studies used traditional machine learning methods such as Support Vector Machine ^4,10^, Random Forest ^5,10,11^, and XGBoost ^14^. Subsequent studies used deep-learning methods that better capture complex or long-range patterns, such as Convolutional Neural Networks (CNNs) ^6^, Long Short-Term Memory (LSTM) ^7,8^, or Transformers ^9,18^. Realizing the importance of considering regulatory and functional relationships between different genes, recent studies also took interaction networks as input and used graph neural networks (GNNs) to identify features that capture gene-gene relationships useful for inferring gene expression levels ^3,14^. Most of these studies also analyzed the relative importance of the different features in the constructed models.

These studies have led to several important observations. First, many features contain non-redundant information about gene expression levels, and thus it is necessary to consider them jointly. Second, some features are related to gene expression levels in a location-specific manner. For example, DNA methylation at the promoter and along the gene body can correlate with gene expression in opposite directions. Therefore, high-resolution data are required to compute features in these different regions separately. Third, all the models constructed so far have not reached the generality required for a universal model. It has been unclear whether this is due to insufficient training samples, inadequate features, or unsuitable modeling methods.

The International Human Epigenome Consortium (IHEC) has recently produced paired epigenomic and transcriptomic data from a large number of diverse human samples using genome-wide sequencing methods that provide base-resolution information. It is an unprecedented resource for modeling transcript levels on a large scale. All IHEC samples and data were quality checked and processed in standardized ways, helping to reduce differences between samples caused by technical confounders.

Using the IHEC data, here we systematically study the feasibility of constructing a universal model of expression levels of transcript isoforms that can be applied across different tissue types. Our study differentiates itself from the previous ones in three main ways. First, we model expression levels of transcript isoforms rather than genes. Since different transcript isoforms of the same gene can be regulated separately and have different expression levels, modeling their expression separately provides substantially more information than grouping all transcript isoforms of the same gene together as a single artificial transcriptional unit. Second, we model the expression levels of transcript isoforms mainly in primary cell and tissue samples. Due to the virtually unlimited supply of cells, cell lines were used in most previous studies to construct gene expression models, but their immortalization and relatively high homogeneity have made it unclear whether the constructed models can be applied to cells from natural tissue contexts. Third, we combine both epigenomic features and multiple types of interaction networks in a single framework to model the expression levels of transcript isoforms. The large amount of data produced by IHEC allows us to define a reasonably large number of epigenomic features for each transcript isoform, while the interaction networks further enable the sharing of information between related genes in the models.

## Results

### Systematic modeling of expression levels of transcript isoforms in 324 samples across 29 human tissue types

To maximize the comparability of modeling results across samples, we selected the subset of samples that have eight types of genome-wide experimental data all available from IHEC to form our core dataset. These eight types of data include expression levels profiled by RNA sequencing (RNA-seq), DNA methylation by whole-genome bisulfite sequencing (WGBS), and histone modifications H3K4me1, H3K4me3, H3K9me3, H3K27ac, H3K27me3, and H3K36me3 by chromatin immunoprecipitation followed by sequencing (ChIP-seq). The resulting core dataset contains 324 samples from 29 tissue types (Figure S1, Supplementary File S1).

To model the expression level of each transcript isoform, we integrated both epigenomic features and interaction networks that describe relationships between different genes, their genomic locations, transcript isoforms, and protein products (Figure S2). Epigenomic features have been shown to correlate with gene expression levels in a location-dependent manner ^1,10,11^. When modeling expression levels of individual transcript isoforms, which often have overlapping genomic loci, it is necessary to define these regions for each transcript isoform separately since the same region can be a promoter of one isoform and an intronic region of another isoform, for example, causing the epigenomic features in this region to be related differently to the expression levels of the two isoforms. Therefore, for each transcript isoform, we defined 21 different regions based on its genomic structure and modeled the relationships between epigenomic signals in these 21 regions and the corresponding expression level of the transcript isoform. These regions cover the upstream, body, and downstream locations of a transcript isoform (Figure S2a, Methods). For each region, we computed the average signal of each of the seven types of epigenomic signals, leading to 21 *×* 7 = 147 epigenomic features per transcript isoform.

In addition to the epigenomic features, different types of interaction networks have also been shown to contain important information about gene expression levels ^14^. Therefore, we defined an integrated network that contains multiple types of interactions, including physical protein-protein interactions (PPIs) and chromatin contacts in three-dimensional (3D) genome structures, together with associations of genes to their transcript isoforms and genomic locations (Figure S2b, Methods). Unlike the epigenomic features, which were all derived from data produced from the corresponding samples, these interactions were either sample-agnostic or obtained from other sources, since sample-matched data were not available (Methods).

We modeled the joint effect of epigenomic features and interaction networks on the expression levels of transcript isoforms using Graph Convolutional Networks (GCNs) ^20^ (Methods). For each sample *i*, we trained a GCN using the whole integrated network and both the epigenomic features and expression levels of transcript isoforms on chromosomes in the training set (Chr1-17 and ChrX). Hyper-parameters were tuned based on modeling performance of transcript isoforms on chromosomes in the validation set (Chr20-22), and we found that 6 hidden layers and 128 hidden nodes per layer to be a good balance between model capacity and generalization (Figure S3, Methods). The resulting model was then applied to both sample *i* and each other sample *j* to infer expression levels of transcript isoforms on chromosomes in the test set (Chr18-19) based on the epigenomic features of the testing sample. We quantified modeling performance by the correlation between the inferred and experimentally measured expression levels across all transcript isoforms on the chromosomes in the test set.

Based on these modeling results, the first question we studied was whether a model learned from sample *i* could infer expression levels of transcript isoforms in another sample *j* as accurately as in sample *i*. In addition, we wanted to know whether the conclusion would be different when samples *i* and *j* come from the same tissue type or not. Putting all our modeling results into these three categories, we found that modeling performance was highest when training and testing a model on data from the same sample (but from different chromosomes, as explained above), followed by having the testing sample different from the training sample but coming from the same tissue type, and lowest when they came from different tissue types (Figure 1a). Similar results were obtained when the rank-based Spearman correlation was used to quantify modeling performance instead of the value-based Pearson correlation (Figure S4), and therefore we only show Pearson correlation values hereafter. Some other biological and technical factors also contributed to the lower performance when applying a model to another sample (Supplementary Text, Figures S5-S6).

**Figure 1:**
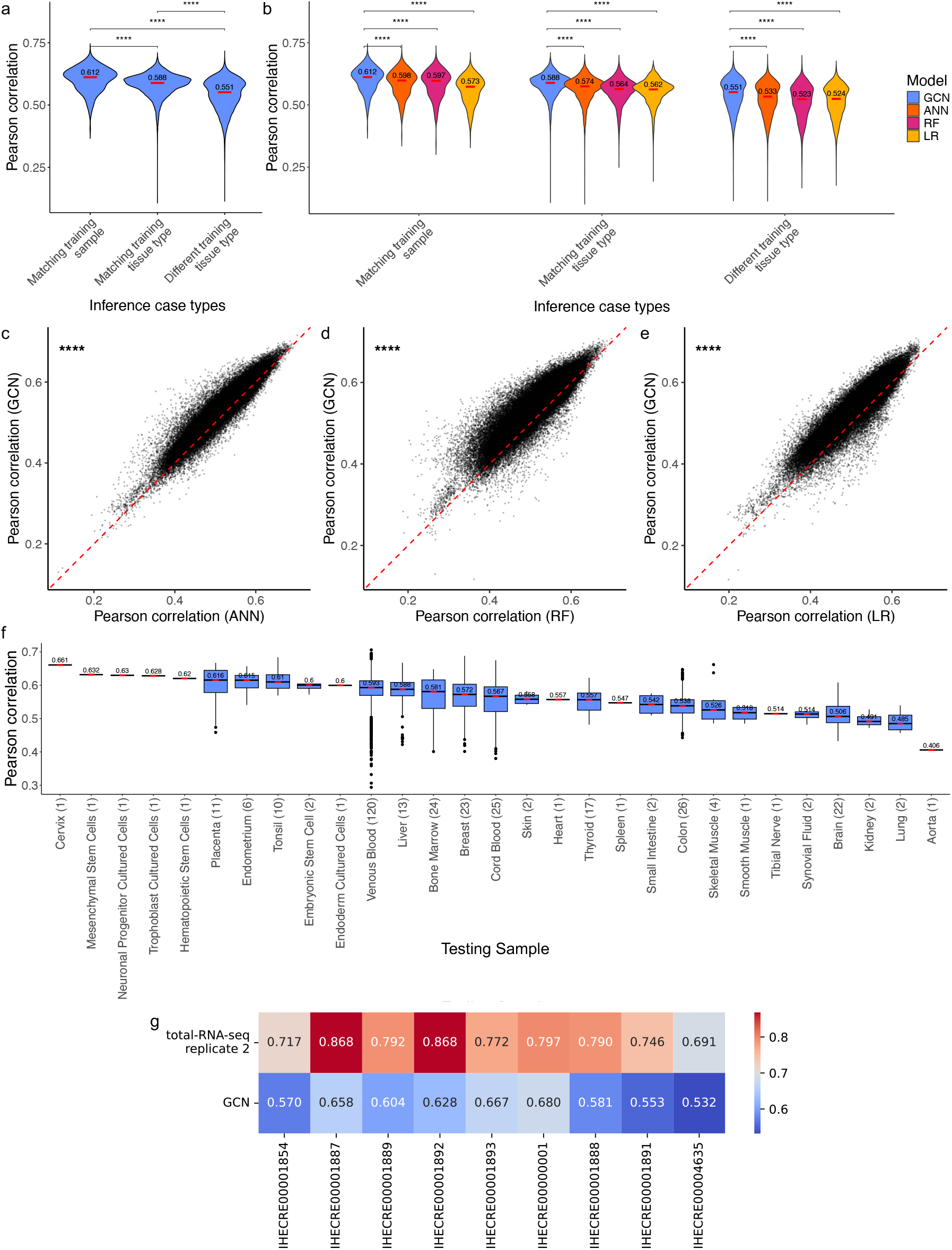
Modeling transcript isoform expression levels using training data from one sample at a time. a, Distribution of modeling performance (quantified by Pearson correlation) when the training and testing samples were either the same, different but from the same tissue type, or from different tissue types. **b,** Comparing modeling performance of Graph Convolutional Network (GCN) with Artificial Neural Network (ANN), Random Forest (RF) and Linear Regression (LR), all three of which do not take the interaction network as input. **c-e,** Comparing modeling performance of GCN with ANN (c), RF (d), and LR (e) in all the individual training-testing sample pairs. **f,** Modeling performance when training and testing models on the same sample, for samples in each tissue type. The number of samples of each tissue type in the whole core dataset is given in parenthesis. **g,** Assessing how close the modeling performance is from the maximum possible performance based on samples with replicated data produced using the total RNA-seq protocol. The two rows show the correlation of transcript isoform expression levels in the test set chromosomes between the two replicates (first row) or between the measured and GCN-inferred levels (second row). In Panels a and b, p-values were calculated using two-sided Wilcoxon rank-sum test. In Panels c-e, p-values were calculated using two-sided Wilcoxon signed-rank test. In all these panels, *:p*<*0.05; **:p*<*0.01; ***:p*<*0.001; ****:p*<*0.0001.

Next, we investigated whether the interaction network contributed to modeling performance, considering that the epigenomic features are already very informative about gene expression levels ^1,10,11^. To do that, we compared performance of our GCN models with three types of classical models that only take the epigenomic features as input but not the interaction network, namely Artificial Neural Network (ANN) (which has the same number of trainable parameters as the GCN), Linear Regression (LR), and Random Forest (RF) (Methods). The results show that the GCN models performed better than these epigenomic features-only alternative models (Figure 1b). These performance gains were consistently observed for many of the individual pairs of training and testing samples (Figure 1c-e). Conversely, we also constructed GCN models with all epigenomic features replaced by random values and found that the modeling performance dropped substantially (Figure S7, compared to Figure 1a). These results show that, by themselves, the epigenomic features contain more information about expression levels of transcript isoforms than the interaction network, but the latter also contains some unique information that complements the former in the models. This dominance of epigenetic features defined around each transcript isoform may also explain why GCN does not drastically outperform ANN. Our goal in this study is to investigate all available data types and the optimal strategy of utilizing them to build a maximal universal model of expression. This is different from our previous work ^14^, where this fine-grained epigenetic feature definition strategy was intentionally excluded, so that the focus was instead to investigate the extent of which interaction networks alone can infer gene expression levels.

To assess the necessity of defining the 21 feature regions per transcript isoform, we constructed alternative models with the 21 regions merged into a small number of regions. These simplified models had reduced performance compared to the original models (Figure S8), thus confirming the usefulness of the 21 regions for defining location-specific epigenomic features, and the unique information present in gene body, promoter, and downstream regions is necessary to comprehensively model expression.

Finally, we investigated whether expression levels of transcript isoforms are easier to infer for certain tissue types when training and testing models on data from the same sample (Figure 1f). Among tissue types with at least 10 samples, placenta models had the highest performance (median Pearson correlation: 0.616), while brain models had the lowest performance (median Pearson correlation: 0.506). This is in part due to the biomaterial type, namely placenta samples were mainly primary cells while brain samples were all primary tissues (Figure S1). The performance difference among tissue types could also be partially caused by the complexity of the composition of cell types, which led to lower performance of the brain models than the thyroid models (Figure 1f), both of which involved only primary tissue samples (Figure S1).

### Estimating distance from maximum possible modeling performance

The above analyses investigated factors that affect the relative performance of different pairs of training and testing samples. Next, we focused on the absolute modeling performance and asked whether it had approached the maximum possible performance. To determine this maximum, we used samples with replicate RNA-seq experiments performed using the same experimental protocol. We set the measured expression levels of the transcript isoforms on the test set chromosomes in one replicate as the target and used their measured expression levels in another replicate as its “predictor” directly without performing any modeling. The resulting correlation between the expression levels of transcript isoforms on the test set chromosomes in these two replicates represents the maximum achievable performance of any model trained on RNA-seq data that are subject to experimental noise and other technical limitations.

In total, there were 16 samples with at least two sets of RNA-seq data produced from the same sample using the same experimental protocol. When both sets of RNA-seq data were measured using the total RNA-seq protocol, their expression levels of transcript isoforms on the test set chromosomes had a correlation ranging from 0.691 to 0.868, which is higher than our modeling performance of 0.532 to 0.680 in these samples (Figure 1g). Similarly, when both sets of RNA-seq data were measured using the mRNA-seq protocol, their transcript isoform expression levels had a correlation ranging from 0.744 to 0.872, which is also higher than our modeling performance of 0.539 to 0.662 (Figure S9). We found that transcript isoform expression level concordance between replicates, while less correlated than gene expression level concordance (Figure S10), remains fairly high considering the expected higher biological variability of transcript isoform expression.

To further assess whether this intrinsic complexity impairs our model, we aggregated transcript-level predictions to the gene level using isoform-length-weighted averaging and obtained a median Pearson correlation of 0.813 against measured gene expression (Figure S11). In addition, when using a per-sample z-score threshold of 1.5 or 2.0 to define expressed transcripts, we classified gene-sample instances with more than three expressed transcripts as challenging and those with three or fewer as non-challenging. We did not observe a reduction in performance for challenging cases relative to non-challenging cases within the same transcript-count group (Figure S12). These analyses indicate that our model remains applicable at both the transcript and gene level, despite the known difficulty of isoform quantification.

We additionally benchmarked our model at the gene level against GC-MERGE ^21^ and Chromoformer ^9^ using the same 7-fold chromosome-stratified evaluation framework. These two methods use Graph Convolutional Network and Transformer architecture respectively, and both utilize epigenetic features and 3D chromatin interaction data, resembling the data types used in our model. Across the three cell lines, our method achieved the highest gene level Pearson correlation among the evaluated methods (Figure S13), indicating that the improvement is preserved after aggregation from isoform-level outputs to gene level expression. We further tested whether chrX and chrY affected the results by repeating the analysis with these chromosomes included or excluded; the correlation changed only minimally, supporting the conclusion that the model generalizes robustly across autosomes and sex chromosomes (Figure S14).

Overall, these results show that our constructed models, trained using data from a single sample at a time, have not reached the maximum possible performance. In the following, we study three potential explanations and explore ways to further improve modeling performance. These explanations include 1) insufficiency of having training data coming from only a single sample, 2) missing other important features, and 3) difficulties caused by specific properties of transcript isoforms.

### Incorporating data from multiple samples into the models

First, we asked whether training a model using data from a single sample was insufficient due to biological and technical variability between samples. We used a meta-learning strategy to approach this question (Figure S15). First, we construct a model using training data from each sample separately, as done above. Then suppose we want to infer transcript isoform expression levels in sample *i* (the “testing sample”), we first select a subset *S* of the constructed single-sample models and apply each of them to infer transcript isoform expression levels in sample *i* separately. Next, we learn a secondary model using these inferred transcript isoform expression levels as features and the actual expression levels in sample *i* as the inference targets. Up to this point, all primary and secondary models are all trained using only chromosomes in the training set. Finally, we apply the secondary model to infer expression levels of transcript isoforms on the left-out test set chromosomes in sample *i*, and compare them with the actual measured transcript isoform expression levels to quantify modeling performance.

We considered three strategies to select the subset *S* of models based on the testing sample *i*:

1. Leave-one-sample-out (LOSO): including all single-sample models trained from samples other than i. This strategy maximizes the amount of training data.
2. Leave-one-tissue-type-out (LOTO): including all single-sample models trained from samples with a tissue type different from that of i. This strategy tests whether the models are general enough that can be applied to an unseen tissue type.
3. Same-tissue-type (ST): including all single-sample models trained from samples with a tissue type the same as that of i. This strategy tests whether including other samples of the same tissue type can improve modeling performance compared to training on data from sample i alone.

From the modeling results, the ST meta-learning strategy led to performance similar to models trained from data of the testing sample (Figure 2a), confirming that the models could generalize across different samples of the same tissue type. Interestingly, the LOSO and LOTO meta-learning strategies led to even higher performance (Figure 2a), and this was observed for almost every individual testing sample (Figures 2b, S16). The superior performance of LOTO is particularly noteworthy, because it shows that without access to data of the testing sample itself or any other samples of that tissue type when training the primary single-sample models, meta-learning is still able to outperform a model trained directly on data of the testing sample (Figure 2b). This result provides crucial support for the possibility of constructing a universal model that can infer expression levels of transcript isoforms across tissue types. To better understand how different samples contributed to the LOTO results, we analyzed their coefficients in the secondary models and observed two major patterns (Figure 2c). First, there were some off-diagonal blocks with strong positive coefficients (black boxes in Figure 2c), which correspond to cases in which data of the contributing samples and testing sample were both produced by the same data production center. However, these cases were not sufficient to explain the generally superior performance of LOTO for almost all testing samples (Figure 2b). Second, some contributing samples were consistently useful when inferring transcript isoform expression levels of other samples (red vertical lines in Figure 2c). We checked all the available metadata of the samples (data source, RNA-seq protocol, etc.) but could not find any of them that can explain these “power contributors”, suggesting that their consistently large coefficients in the secondary models reflect important biological information they contain rather than technical confounders.

**Figure 2:**
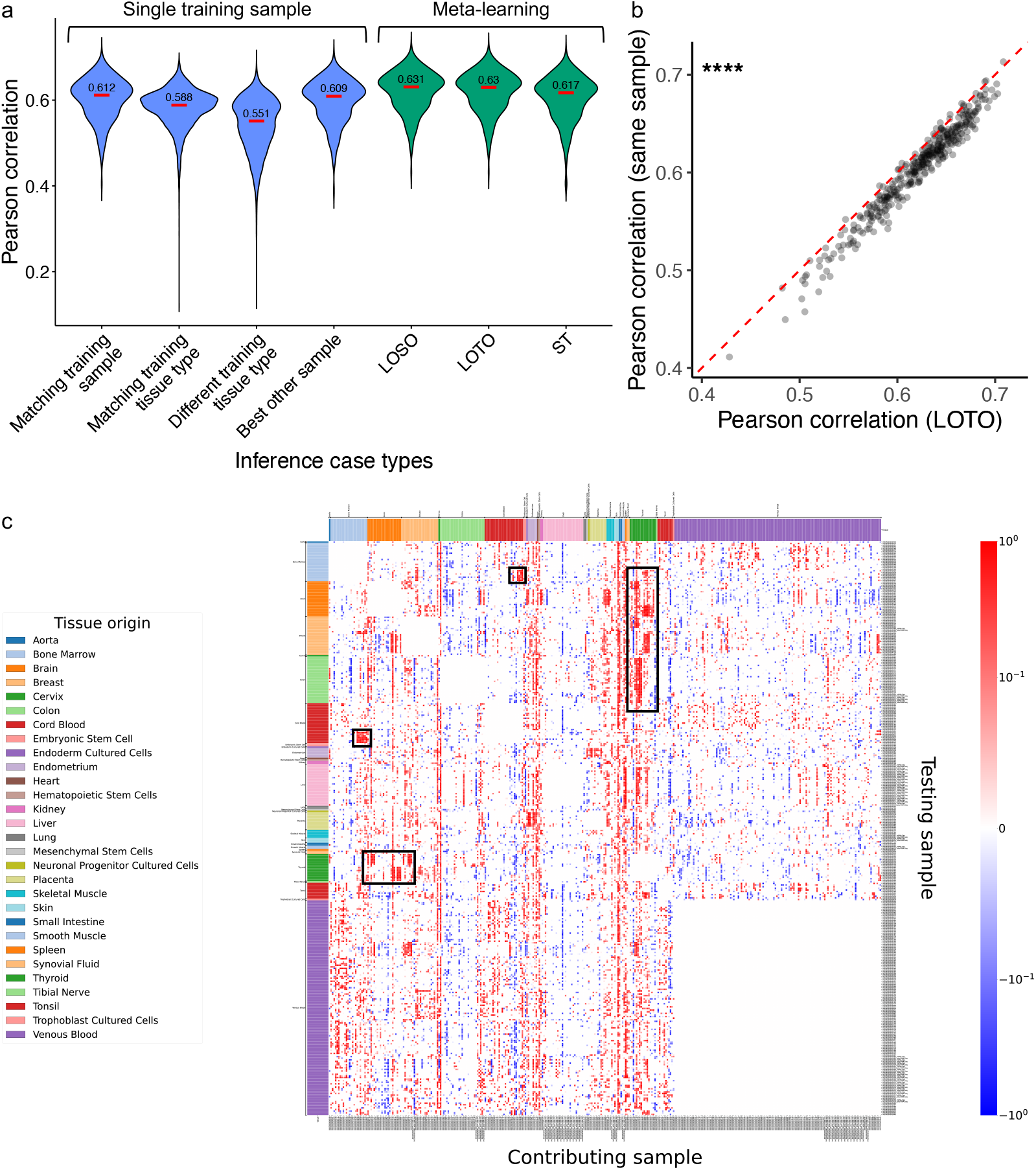
Modeling transcript isoform expression levels using meta-learning of models constructed from multiple training samples. a, Comparing modeling performance of different single-sample and meta-learning model construction strategies. The first three distributions are copied from Figure 1a for easy comparison. **b,** Comparing modeling performance of single-sample models trained on data from the testing sample with secondary models produced using the LOTO meta-learning strategy. **c.** Secondary model coefficients of the LOTO meta-learning strategy. Each row represents a testing sample the transcript isoform expression levels of which are to be inferred by the meta-learning procedure. Each column represents a sample contributing to the secondary model. In the heatmap, the color of an entry indicates the coefficient of the contributing sample in the secondary model constructed for the testing sample, which can be either positive (red) or negative (blue). Since each sample does not contribute to secondary models of samples of the same tissue type, all entries in the diagonal blocks are zero. In Panel b, p-value was calculated using two-sided Wilcoxon signed-rank test. *:p*<*0.05; **:p*<*0.01; ***:p*<*0.001; ****:p*<*0.0001.

In view of these results, we checked whether the model of a single power contributor could perform as well as the secondary model obtained using the LOTO or LOSO meta-learning strategy. Specifically, for each testing sample *i*, we selected another sample *j* that produced the highest single-sample modeling performance. These models (“Best other sample” in Figure 2a) already approached the performance of training a model using data from the testing sample *i* itself (“Matching training sample” in Figure 2a). Interestingly, only 69% of these best other samples came from the same tissue type as the sample *i*, showing that in the remaining around 30% of the cases, training data from another tissue type were even more useful. However, these best other sample models still could not reach the performance of the LOTO models (Figure 2a), showing that it is advantageous to combine data from multiple samples using the meta-learning strategy.

### Incorporating additional features into the models

Next, we studied whether modeling performance can be improved by including additional features. Since chromatin accessibility is a commonly used proxy of functional potential ^22^, we compared our original single-sample GCN models with ones that also have chromatin accessibility features derived from ATAC-seq or DNase-seq data (or both) on samples with such data available (Methods). Judging by the correlation between the inferred and experimentally measured transcript isoform expression levels, adding chromatin accessibility features did not consistently improve modeling performance (Figure 3a). To investigate the reason, we analyzed the importance of different features in the models using saliency and feature value permutation (Methods). Saliency measures the dependency of a model output on an input feature, while feature value permutation determines the importance of a feature by the increase of modeling error if the values of the feature are randomly permuted. From the saliency results, the model outputs had some dependency on the ATAC-seq features in the first exon and the first intron (Figures 3b, S17a) but not so much on the DNase-seq features in general (Figures 3b, S17b). This analysis also revealed that the model outputs depended most on i) DNA methylation in gene body regions and around the transcription start site (TSS), ii) H3K36me3 in gene body regions, especially the internal introns, and iii) ATAC-seq features in the first exon (Figure 3b). From the feature value permutation results, the DNase-seq features were not particularly important in the models while ATAC-seq features at several regions were more important when DNase-seq features were not supplied as input (Figures 3c, S17c-d). Interestingly, DNA methylation features were considered important by the saliency method but not feature value permutation, while H3K36me3 features at internal introns were usually considered important by both methods (Figures 3b-c, S17a-d). This may indicate that while DNA methylation features are useful for inferring transcript isoform expression levels, the information that they contain are not as unique as the H3K36me3 features. Overall, these results show that some chromatin accessibility features contain useful information and they contributed to the models, but given the presence of other strong features such as those derived from DNA methylation and H3K36me3, the gain in modeling performance caused by the chromatin accessibility features was small.

**Figure 3:**
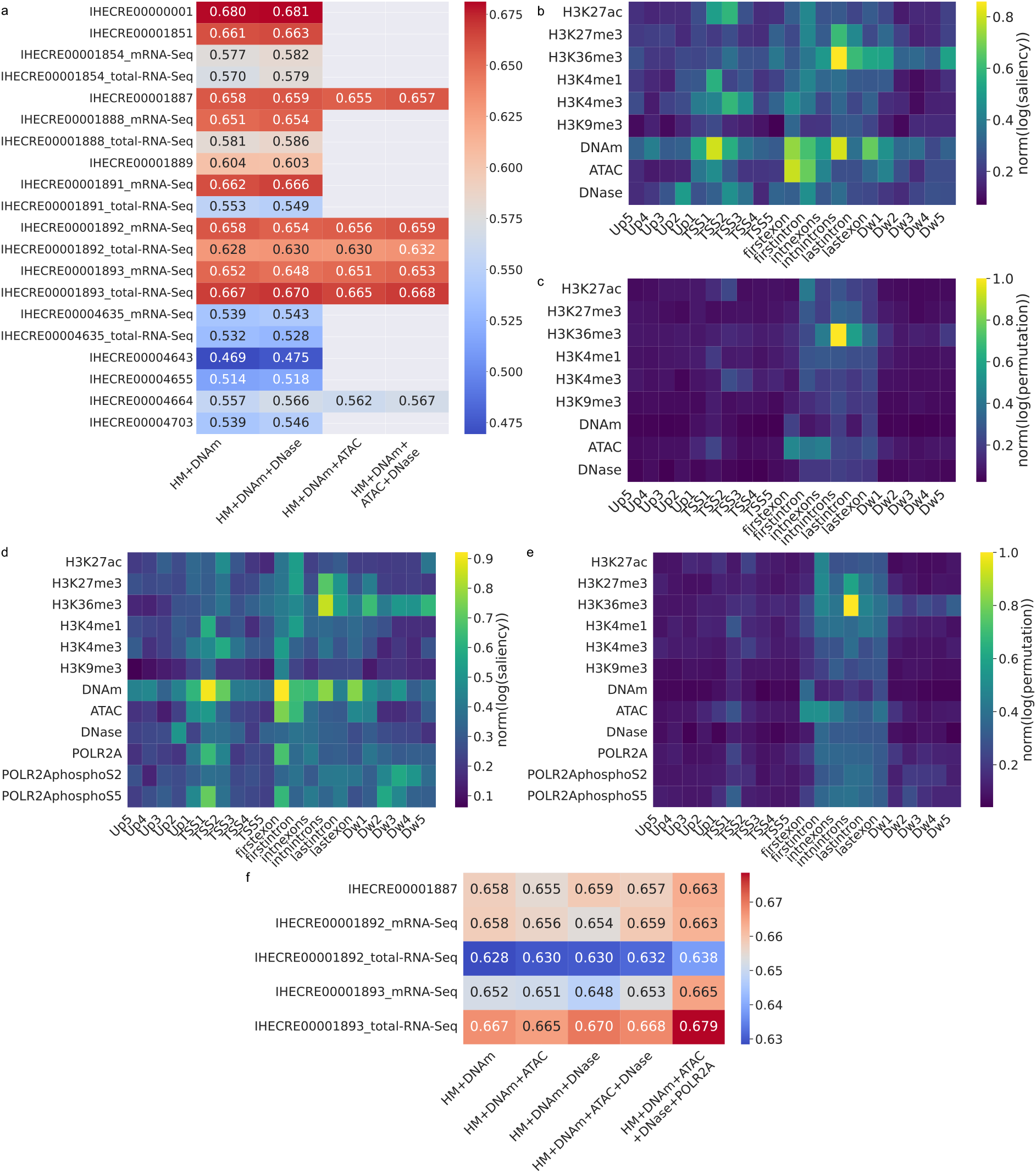
Effects of incorporating additional features into the single-sample models. a, Comparing modeling performance of the original models, which contain features derived from 6 types of histone modification and DNA methylation (“HM+DNAm”), with new models that also contain chromatin accessibility features from either DNase-seq, ATAC-seq, or both. **b-e,** Importance scores of features derived from the original seven types of epigenomic features plus those derived from chromatin accessibility data (ATAC-seq and DNase-seq) only (b-c) or also with Pol II features (d-e). Importance is quantified by saliency (b and d) or feature value permutation (c and e). **f,** Comparing modeling performance with and without POLR2A features.

Since our core dataset contained only data for 6 types of histone modification, we tested whether incorporating additional types of histone modification could improve modeling performance. However, for samples that had such data available, these extra features only marginally improved modeling performance, even when 21 additional types of histone modification were added (Figure S18). This indicates that our original set of histone modification and DNA methylation features already contained most of the same information about transcript isoform expression levels.

For three of the samples, we also had ChIP-seq data for RNA polymerase II (POLR2A subunit) binding, including both the unmodified form and the forms with phosphorylated Serine 2 and Serine 5 (phosphoS2 and phosphoS5), which are detected primarily in coding regions and promoter regions, respectively ^23^. We expected these three types of POLR2A features to be strong indicators of transcript isoform expression levels. Indeed, some model outputs depended on these features and overall modeling performance had some improvements, although these features were not considered most important as compared to the other features (Figures 3d-f). Relatively, the unphosphorylated and phosphoS5 forms had higher saliency scores than the phosphoS2 form (Figure 3d).

As a key player of transcriptional regulation, we explored whether transcription factor (TF) binding features could further improve modeling performance (Methods). In general, adding TF features did not lead to clear improvements of modeling performance even when binding data for *>*700 TFs were added (Figure S19), suggesting that they did not bring much new information to the models.

Overall, these results show that adding extra chromatin accessibility, histone modification, or protein binding data led to some improvements of modeling performance, but not as consistently as incorporating data of the 7 core feature types from more samples using the meta-learning strategy.

### Differences in modeling performance among transcript isoforms

Finally, we investigated properties of transcript isoforms that affect modeling performance. First, we asked how the variability of expression levels of a transcript isoform across different samples affects its modeling performance. We found that transcript isoforms with the lowest standard deviation of expression across samples had the lowest modeling performance (Figure 4a), likely because these small differences are more difficult to model and more affected by noise. On the other hand, after normalizing by mean expression across samples, the resulting coefficient of variation (CV) displays the opposite trend, where transcript isoforms with a higher CV had worse modeling performance (Figure 4b). Some of these transcript isoforms may be expressed only in some specific tissue types of samples and their epigenomic features may show some unique patterns that the models did not fully capture.

**Figure 4:**
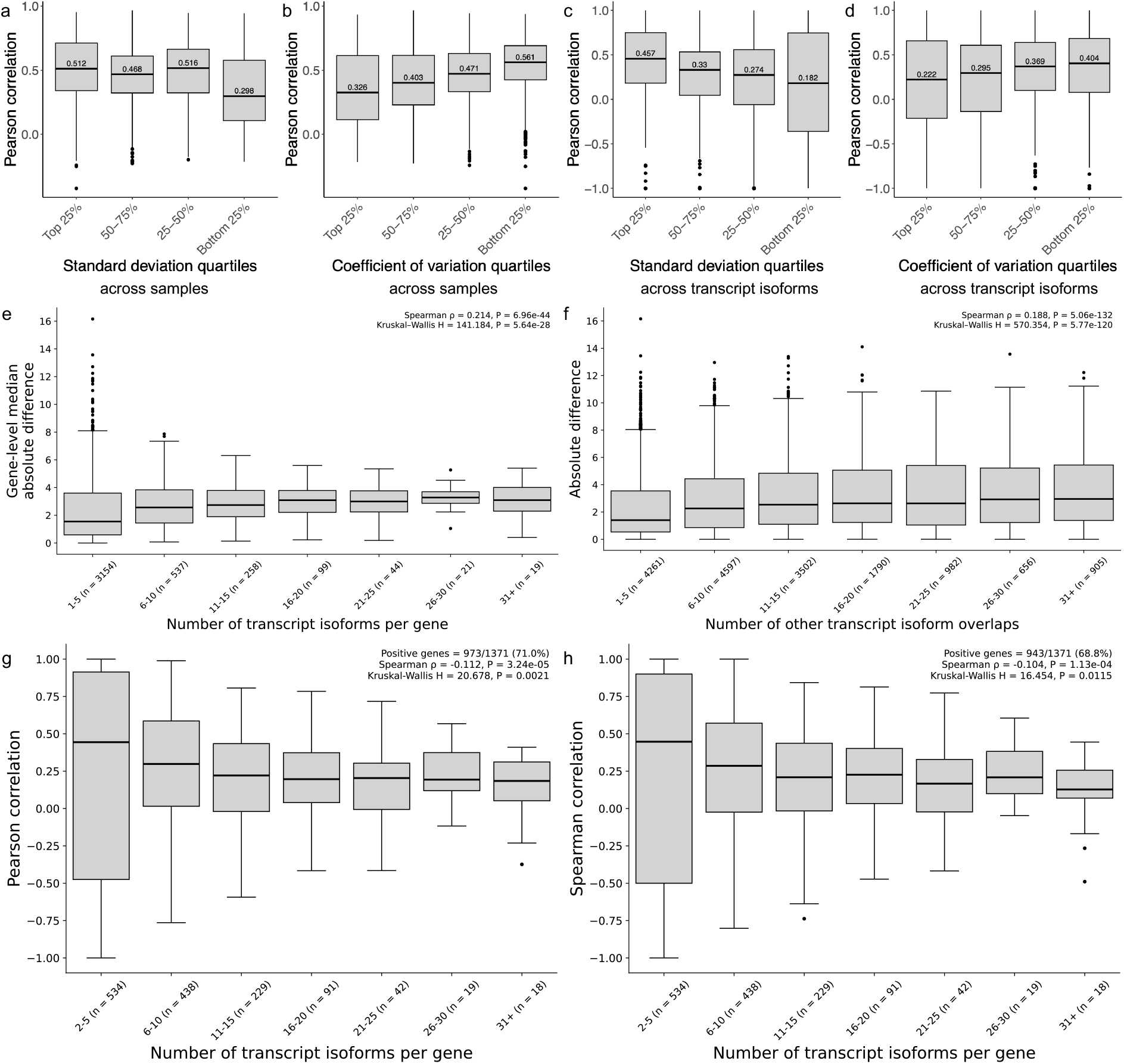
Properties of transcript isoforms that affect their modeling performance a-d, Modeling performance of transcript isoforms with different expression level variability across samples (a-b) or across the transcript isoforms of the parent gene (c-d), quantified by standard deviation (a and c) or coefficient of variation (b and d). Data shown in Panels c and d are from sample IHECRE00000001. **e-f,** Modeling performance of transcript isoforms with different number of transcript isoforms of its parent gene and binned for visualization (e) or different number of other isoforms overlapping it and binned for visualization (f). Here modeling performance is quantified by absolute difference between inferred and measured expression levels since for genes with a small number of transcript isoforms, Pearson correlation is not informative. **g-h,** Comparing inferred and actual expression levels of different transcript isoforms of each gene binned for visualization, quantified by Pearson correlation (g) and Spearman correlation (h).

Next, we asked how the variability of expression of different transcript isoforms of the same gene affect their modeling performance. Similar to the results above, we found that modeling performance was lower for genes with a small standard deviation among expression levels of its transcript isoforms (Figure 4c) or a higher CV among them (Figure 4d). We speculated that the latter is highly related to the number of transcript isoforms a gene has, namely a gene with more transcript isoforms tends to have a higher CV of their expression and also lower modeling performance in general due to the complexity. Indeed, we found that modeling error tended to be higher for genes with more transcript isoforms (Figure 4e).

In addition to the number of transcript isoforms that a gene has, how much the exons of the different isoforms overlap each other also contributes to the complexity of gene regulation and difficulty in modeling. Biologically, these different isoforms share common epigenomic signals and common proximity to regulatory elements and TF binding, and thus it is difficult to control their expression levels separately. Technically, sequencing reads produced from the overlapping regions cannot be uniquely assigned to one transcript isoform, leading to difficulties in determining the precise expression level of each isoform. Consistent with these complexities, we found that transcript isoforms that overlapped with a larger number of other isoforms tended to have higher modeling errors of their expression levels (Figure 4f).

Considering the effects of isoform complexity on modeling performance described above, we also asked how well our models were able to infer expression levels of different isoforms of the same gene. As opposed to the other analyses presented above, in which we evaluated transcript isoforms from different genes together, this time we took transcript isoforms of one gene at a time and computed the correlation between inferred and measured expression levels of the transcript isoforms of this gene only. Then putting these correlation values from different genes into groups based on number of transcript isoforms per gene, we found that i) the resulting median correlation values were mostly positive, and ii) modeling performance did not clearly decrease with an increasing number of transcript isoforms (Figure 4g-h). After annotating each bin with the number of genes it contained and excluding the lowest isoform-count bin, in which correlation estimates are inherently unstable because they are computed from very few isoforms per gene, the median within-gene correlation remained generally positive and the rank-based Spearman analysis showed the same overall trend as Pearson correlation (Figure S20). Therefore, our models showed capability of telling which isoform of a gene has a higher expression level than another isoform regardless of the number of isoforms the gene has, but having more transcript isoforms per gene tended to make it more difficult for our models to determine their expression levels relative to other genes. To assess whether transcript isoforms from the same gene are sufficiently similar in epigenetic feature space to justify collapsing them into a single representation, we analyzed pairwise differences across 7 epigenetic marks and 21 genomic regions per transcript using PCA under two complementary settings. In setting 1, same-gene isoform pairs were compared against pairs drawn from different genes; in setting 2, same-gene pairs with high versus low expression were compared with expression-stratified pairs from different genes. Although PC1 showed partial separation between same-gene and different-gene isoform pairs, substantial overlap remained between the distributions, with even greater overlap along PC2. These results indicate that same-gene isoform pairs are not consistently more similar than different-gene pairs in epigenetic feature space, suggesting that isoforms retain considerable epigenetic heterogeneity and should not be collapsed solely based on their shared gene origin (Figure S21). We next directly examined whether epigenetic divergence between same-gene isoforms was associated with differences in their expression. For each sample, we calculated the absolute expression difference between isoform pairs and compared it with the corresponding divergence in their 147-feature epigenetic profiles. Pairs with high expression differences showed significantly greater Euclidean distances than pairs with low expression differences, indicating that more transcriptionally divergent isoforms tend to have more divergent epigenetic features (Figure S22a). Consistently, high-expression-difference pairs showed lower cosine similarity between their epigenetic profiles (Figure S22b). However, substantial overlap remained between the two groups, with many pairs retaining highly similar epigenetic features despite large expression differences. We further examined epigenetic divergence across individual chromatin marks and genomic positions. Differences between lowand high-expression-difference pairs were observed across multiple marks and genomic regions (Figure S22c,d), indicating that the observed epigenetic divergence is not restricted to a small subset of features. Together, these analyses show that same-gene transcript isoforms can exhibit distinct epigenetic feature representations and do not support merging isoforms solely on the assumption that their epigenetic features are redundant.

## Discussion

In this study, taking advantage of the large amount of paired epigenomic and transcriptomic data produced by IHEC, we quantitatively modeled the expression levels of transcript isoforms by integrating epigenomic features and different types of biological interactions. In general, the expression levels inferred by our quantitative models correlate with the actual expression levels fairly well as evaluated by left-out data from test set chromosomes. When training models using data from a single sample alone, we found that modeling performance was affected by differences between samples and tissue types, properties of transcript isoforms, and other biological and technical factors such as disease status and experimental protocols.

To explore the possibility of forming a universal model that can infer transcript isoform expression levels accurately in any tissue type, we considered a meta-learning strategy that combines information from multiple samples. We found that incorporating data from multiple training samples could improve modeling performance in general. Importantly, results of our Leave-one-tissue-type-out (LOTO) strategy showed that even without training data from the target tissue type when constructing the primary single-sample models, the resulting secondary meta-learning models still consistently outperformed models trained on data from the testing sample itself. This suggests that it is possible to capture general quantitative relationships between transcript isoform expression levels and epigenomic and network features that apply across tissue types.

We also investigated the possibility of improving model performance by incorporating additional features. In general, including additional features derived from chromatin accessibility, extra types of histone modification, POLR2A binding, or TF binding only led to some small improvements of model performance, at a level less pronounced than including training data from more samples. This suggests that for the purpose of modeling the expression levels of transcript isoforms, it is more advantageous to include more samples than profile more types of epigenomic signals.

Using the meta-learning strategy, the inferred expression levels of our models had a median correlation of 0.63 with the actual measured levels. In comparison, the median correlation between two replicated RNA-seq experiments was 0.79. We speculate that this performance gap is due to post-transcriptional regulation that our models were not able to capture. In the future, it would be useful to incorporate features or interaction networks about post-transcriptional regulation into the models, or replace RNA-seq data by data that reflect rate of nascent transcript production when training and testing the models. Both approaches are expected to narrow the gap between the performance of the models and the maximum possible performance.

Recently, Transformers using DNA sequences as input features ^9,18,19,24,25^, specifically framed as a track coverage modeling task, have gained popularity. These methods have fundamentally different input features than our method, which takes epigenetic features and an interaction network as input. A key distinguishing factor between our method and DNA sequence-based Transformer models is their inability to be applied to a new sample if the DNA sequence of the new sample is unknown or is identical to a training sample (e.g. from another cell type of the same individual). Thus these models cannot be used for the type of cross-sample predictions heavily demonstrated for generalizability and cross-tissue ensemble in this work. An advantage of these models is that they can predict effects of DNA mutations but this is generally difficult for our model due to the lack of large scale epigenomic datasets for targeted mutations and their subsequent genome-wide epigenetic change.

Although most of the samples in our core dataset were primary tissues or primary cells, a small proportion of the samples were immortalized cell lines. We found that the expression levels of transcript isoforms in cell lines were easier to infer regardless of whether the model was trained on data from a cell line or not (Figure S5a). Compared to cell lines, tissue samples are more complex because they have a mixture of cell types. As a result, what our models captured were relationships between cell-averaged epigenomic features and cell-averaged transcript isoform expression levels. It is unlikely that such cell type mixture rendered our models unable to reach the maximum possible performance because even the cell line models were not able to achieve it. However, we expect that having an estimate of the cell type composition in each sample can help improve our models.

If sparsity issues can be overcome, it would also be useful to test and refine our models using single-cell multi-modal data that have both gene expression and epigenomic features profiled in the same cell, especially paired RNA–ATAC profiles, to reduce confounding from bulk cell-type composition and to enable cell-type-specific regulatory inference. However, the sparsity of single-cell assays and the limited set of epigenetic modalities that can currently be profiled from the same cell remain important practical constraints. In addition, technical issues of current single-cell RNA-seq data affect the reliability of expression measurements, in particular dropouts affecting low expression gene quantification, and individual transcript isoform quantification being uncommon remain as major challenges to be resolved before reliable single-cell datasets can be utilized in our methods. There are also existing methods that infer gene regulatory networks (GRNs) from single-cell data (scRNA-seq alone or scMultiome), for example by combining co-expression, sequence motifs, and prior knowledge. However, how these inferred GRNs should be used to model gene expression levels in single cells requires careful attention to avoid information leakage, specifically when using gene expression information prior from scRNA-seq to infer the GRN, and subsequently using it to model expression levels.

To assess the potential technical biases of datasets collected from various sequencing projects despite rigorous QC steps and standardized processing pipelines by IHEC, we have systematically compared how different technical factors affect modeling performance. These technical factors, while contributing to performance shifts of Pearson correlation between predicted and measured transcript expression levels, never exceeded 0.05 for each factor. Similarly, these technical factors cannot clearly separate samples with large coefficients in the meta-learning LOTO tests. Specifically, when focusing on the RNA-seq protocol used as label source, we found that when training and test data involve the same RNA-seq protocol, the models based on total RNA-seq performed slightly better than the models based on mRNA-seq with a median correlation of 0.570 and 0.555 respectively (Figure S5c), potentially due to the expected inability or bias in capture of transcripts without a poly-A tail such as those from some non-coding and histone genes. On the other hand, applying models trained from mRNA-seq data to predict expression levels in total RNA-seq data did not cause performance degradation, with a median correlation of 0.571, suggesting that the technical differences between the two RNA-seq protocols did not necessarily make the models learned clearly less generalizable.

In this work, we have used some epigenomic and network features that can causally affect transcript isoform expression levels; however, some features included can also simply numerically infer transcriptional activities without playing a causal role. For example, protein-protein interactions can be involved in transcriptional regulation by defining specific transcription factor complexes, but they can also be downstream effects of transcription, namely proteins that physically interact must be expressed at the same time (at the protein level, which is partially, but not fully, reflected by transcript levels). Therefore, the models constructed in this study should not be interpreted as ones that can mechanistically infer transcript isoform expression levels.

In addition to modeling transcript isoform expression levels in a single sample, another related problem is modeling differential expression between two samples or two conditions, such as disease and non-disease states. A universal model of transcript isoform expression levels in a single sample can be used to infer differential expression between two samples, but before such a universal model becomes available, modeling differential expression could be a separate problem with unique features and modeling methods ^2,8,11^.

## Methods

### Constructing the core dataset

We selected 324 samples with genome-wide measurements of transcript expression, six types of histone modification (H3K4me1, H3K4me3, H3K9me3, H3K27ac, H3K27me3, and H3K36me3), and DNA methylation all available to form our core dataset (Supplementary File S1). All data were downloaded from the IHEC EpiATLAS Data Portal (https://ihec-epigenomes.org/epiatlas/data/) with a data cutoff date of August 29, 2023. This dataset is the newly expanded successor of the previously published IHEC resource ^26^, and our current analysis uses the updated IHEC collection ^27^, which fully subsumes all samples and results from the earlier published dataset. Accordingly, our conclusions reflect performance across both the original and newly generated IHEC resources, which includes data generated from AMED-CREST, BLUEPRINT, CEEHRC, DEEP, ENCODE, EpiHK and NIH Roadmap.

Specific details of different types of data are provided below.

### Transcript isoform expression levels from RNA-seq data

We obtained the quantified expression level of each transcript isoform in each of the 324 samples from the IHEC portal. The data were produced using two different RNA-seq protocols, namely the mRNA-seq protocol that performs poly-A enrichment and the total RNA-seq protocol that performs ribosomal RNA depletion without poly-A enrichment. For 34 samples with both types of data produced, we treated each of them as two separate samples with identical epigenomic features in the modeling tasks, leading to 324 + 34 = 358 “target samples” in total.

RNA-seq data were processed by IHEC using a standardized Nextflow pipeline. For samples with replicates, data from the different replicates were combined before producing the final transcript isoform expression levels. Transcript isoform annotations were obtained from GENCODE ^28^ (v29), which is based on the hg38 reference human genome. We took the transcripts per million (TPM) values of each transcript isoform, added a pseudocount of 1e-3 to it, and then performed log-2 transformation of the resulting value. In total, we analyzed the complete transcriptome of 205,759 transcript isoforms from 58,121 protein-coding and non-coding genes, all of which were quantified by IHEC’s standardized pipeline in each of the 324 samples, with expression values that could be zero or non-zero.

### Locations of epigenomic features

For each transcript isoform, we defined 21 regions for computing the epigenomic features (Figure S2a):

- Promoter (5 regions): the 2kb region immediately upstream of the TSS is divided into five 400bp regions labeled as Up1 (closest to TSS) to Up5 (farthest from TSS)
- TSS-proximal (5 regions): the 2kb region immediately downstream of the TSS is divided into five 400bp regions labeled as TSS1 (closest to TSS) to TSS5 (farthest from TSS)
- Downstream (5 regions): the 2kb region immediately downstream of the transcription end site (TES) is divided into five 400bp regions labeled as Dw1 (closest to TES) to Dw5 (farthest from TES)
- Gene body (6 regions): six regions are defined, including first exon, internal exons, last exon, first intron, internal introns, and last intron. Internal exons are defined as the exon(s) excluding the first and last exons. If there are multiple internal exons, they are concatenated into a single virtual contiguous region. Internal introns are defined in the same way.

This windowing approach was adopted from prior epigenomic modeling studies to capture localized regulatory signals that can contain enhancers and splicing signals while maintaining manageable feature dimensionality ^4,10,11,29,30^. Defining multiple contiguous windows enables the model to resolve positionspecific patterns, such as promoter-proximal versus gene body histone modifications, which are known to differentially correlate with gene and transcript expression levels.

In order to have meaningful epigenomic features for all the regions, a transcript isoform needs to have at least 3 introns (i.e., at least 4 exons). For other transcript isoforms, we filled in the dummy value of-1 for features that could not be defined.

For each transcript isoform, its 15 promoter, TSS-proximal, and downstream regions are all nonoverlapping, and its 6 gene body regions are all non-overlapping. On the other hand, the TSS-proximal regions can overlap with the gene body regions (always overlapping with first exon, and sometimes overlapping with first intron or other gene body regions).

### DNA methylation features from WGBS data

WGBS data were processed by IHEC using a standardized pipeline modified from gemBS ^31^. We obtained beta values of each CpG site (i.e., proportion of reads with an unconverted cytosine among all aligned reads) in each sample from the IHEC portal. For each of the 21 regions of a transcript isoform, we defined its DNA methylation feature as follows. First, for each CpG site *i* within this region with aligned reads in the processed and filtered data, we record the number of them that support the site to be methylated, *m_i_*, and the number of them that support the site to be unmethylated, *u_i_*. We then aggregated all these numbers to become the final methylation level of the whole region as 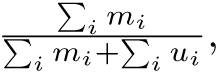 where the summation is taken over all CpG sites in the region. For regions with no aligned reads, we set the feature value to-1.

### Histone modification features from ChIP-seq data

ChIP-seq data were processed by IHEC using a standardized pipeline modified from the ENCODE pipeline (https://www.encodeproject.org/chip-seq/histone/). We obtained fold-change-over-control values for each genomic position from the IHEC portal, and calculated an average score for each of the 21 regions of a transcript isoform in each sample. We added a pseudocount of 1e-9 to each value, performed log-2 transformation of it, and then shifted the value by deducting the log-2 transformed pseudocount from it. Finally, we z-scaled the log-2 transformed value and clipped it to the range of-2.5 to 2.5, and shifted the value to the range of 0 to 5. For regions with no data, we set the feature value to-1.

### Additional features

For a subset of 14 samples originating from the ENCODE project, we collected DNase I hypersensitivity assay and/or ATAC-seq data (14 samples with DNase I hypersensitivity assay data, 4 samples with ATAC-seq data, 4 samples with both) and constructed the 21 regions of each transcript isoform using the same method described above for histone modification ChIP-seq data. Out of these 14 samples, 3 also had POLR2A ChIP-seq data (POLR2A, POLR2AphosphoS2 and POLR2AphosphoS5), which we again constructed the 21 regions of each transcript isoform using the method for histone modification ChIP-seq data. For these models, we only reported their performance when testing on the samples that they were trained on (based on the left-out testing chromosomes).

Additionally, for a subset of 9 samples originating from the ENCODE project, we collected additional histone modification ChIP-seq data and constructed the 21 regions of each transcript isoform using the same method outlined above for histone modification ChIP-seq data. We further selected 5 samples with large amounts of transcription factor ChIP-seq data from ENCODE and generated extra TF feature sets. These include: a) 1 feature (promoter region only, which is the average feature score in the region from 1,000 bp upstream to 500 bp downstream of TSS), b) 3 features (promoter, exon and intron regions) and c) 5 features (promoter, exon, intron, full gene body and downstream (1000 bp downstream of TES) regions).

Notably, these additional profiles were obtained from ENCODE rather than IHEC, providing us with additional ATAC-seq, DNase-seq, TF ChIP-seq and extra histone modification datasets, instead of the original 6 histone marks and DNA methylation epigenetic features, allowing us to assess transfer across datasets and to test whether performance gains persist when the model is exposed to an independent source of epigenomic measurements. This provides both a biological and technical validation of generalization beyond the original cohort.

### Defining the integrated interaction network

Our integrated interaction network consisted of three types of nodes, namely i) genes and their protein products (denoted as “g”); ii) transcript isoforms of the genes (denoted as “t”), and iii) non-overlapping 50kb regions that tile the whole genome termed genomic bins (denoted as “b”). The nodes were connected by four types of unweighted edges: a) g-t interactions that connect each gene to all of its transcript isoforms; b) g-b interactions that connect each gene to all genomic bins that overlap it; c) g-g interactions that connect genes whose protein products have physical protein-protein interactions (PPIs); and d) b-b interactions that connect genomic bins with significant contacts in 3D genome structures. In total, our network contained 58,121 g nodes, 205,759 t nodes, 60,630 b nodes, 411,518 g-t interactions, 189,778 g-b interactions, 1,560,111 g-g interactions, and 39,704,973 b-b interactions.

Below we provide additional details about the g-g and b-b interactions.

### Protein-protein interactions (g-g)

We obtained PPIs from BioGRID ^32^ (v3.4.162). We considered only interactions specified to be physical interactions and excluded other types of interactions such as genetic interactions or co-expression information.

### Chromatin contacts in 3D genome structure (b-b)

We collected Hi-C data produced from 10 cell lines from two different sources. For six cell lines, including HCT-116, HMEC, HUVEC, IMR90, K562 and NHEK, we downloaded the raw reads from the Sequence Read Archive (SRA) ^33^ (project accession numbers SRP118999 and SRP050102) and aligned reads in each library to the hg38 reference human genome using the Burrows-Wheeler Aligner (BWA) ^34^ (v0.7.17r1188). We retained only sequencing reads with a mapping quality score of 30 or more and used Juicer ^35^ (v1.6.0) to generate a merged.hic file for each sample across all its sequencing libraries.

For another four cell lines, including GM12878, H1-hESC, HeLa-S3, and HepG2, we obtained the processed Hi-C datasets (aligned to the hg38 reference human genome) directly from 4D Nucleome ^36^ (accession numbers 4DNFI1OUWFSC, 4DNFIQYQWPF5, 4DNFICEGAHRC, and 4DNFICSTCJQZ, respectively).

For each of the 10 cell lines, we produced the contact matrix between 50kb genomic bins using juicer-tools with KR normalization. Following a previous work ^14^, we called significant interactions with FitHiC2 ^37^ (v2.0.7) using the q-value threshold of 0.005 and set the “chromosome region” parameter to “All”, to consider both inter-and intra-chromosomal Hi-C interactions. These interactions served as the bin-bin interactions in our integrated biological network. If a pair of genomic bins have a significant Hi-C interaction in multiple cell lines, we kept only one edge between them.

### Construction of transcript isoform expression models

We treated our integrated interaction network as an undirected network by having a reverse edge for every edge defined. We also added a self-loop to each node. These nodes formed the first layer of the GCN model. Each of the input transcript isoform (“t”) nodes was associated with 21 *× f* features, where by default *f* = 7 for the seven types of epigenomic features when the additional features (chromatin accessibility, POLR2A, extra HMs, or TFs) were not included. For the gene (“g”) and genomic bin (“b”) nodes, the same number of features were also defined, all filled with the dummy value of-1. We then stacked 6 hidden graph convolutional layers with 128 hidden neurons in each layer, and a dropout layer applied before each hidden graph convolutional layer, followed by an output layer that reports the inferred expression level of each transcript isoform. This configuration balances expressive capacity with overfitting risk. The depth and width were guided by prior work in GCN architectures for biological networks, which indicates that these numbers of neurons and layers are sufficient in effectively capturing complex regulatory relationships without excessive parameterization ^21^. We selected the GCN architecture by systematically varying hidden-layer width (32, 64, 128, 256 and 512 nodes) and depth (1 to 9 layers) based on validation-set Pearson correlation. Wider layers generally allowed useful performance to persist across greater depths, while excessive depth led to diminishing returns. In particular, 128 nodes maintained near-optimal validation performance across 1 to 6 hidden layers, after which performance declined. We therefore selected 128 nodes per layer and 6 hidden layers as a good balance between model capacity and generalization. Notably, 256 and 512 nodes substantially increased training performance without improving validation performance, suggesting increased risk of overfitting. The independent test results further supported this choice. We used the rectified linear unit (ReLU) as our non-linear activation function at each layer besides the output layer. We implemented our GCN models using the “GraphConv” module of Deep Graph Library (DGL) ^38^ (v0.6.1). PyTorch (v1.8.1) was used as the back-end of DGL with CUDA 11.1 support.

We used minimum squared error (MSE) as the loss function to quantify the difference between the inferred and actual transcript isoform expression levels. We trained our models using the Adam optimizer ^39^ with a learning rate of 0.0005 and a weight decay of 0.0005, and trained our models with 5000 epochs and applying a dropout of 0.45. We trained our models using either NVIDIA A40 or NVIDIA GeForce RTX 3090 GPUs, depending on the GPU memory required for a model.

We divided transcript isoforms on different chromosomes into a training set (Chr1-17 and ChrX), a validation set for hyper-parameter tuning (Chr20-22), and a left-out test set for model evaluation (Chr18-19). In this setting, around 85% of the transcript isoforms were used for training, and the longest chromosomes were used to define the training set to maximize the amount of training data. This chromosome-based splitting strategy ensures that transcripts in the training, validation, and test sets are genomically independent, preventing information leakage from transcript isoforms of the same gene and providing a more stringent test of model generalization. Chromosome stratification was selected over random transcript sampling to control for potential cis-regulatory correlations within chromosomes, following prior studies using chromosome-stratified evaluations in gene expression modeling ^14^. Specifically, this is to prevent the scenario where a transcript isoform is in the training set and another transcript isoform from the same gene is in the test set, in which case the model may simply remember the specific epigenetic features and expression level of the former and use them to infer the expression level of the latter.

All modeling performance reported in this manuscript, including performance of models learned from single samples and performance of the meta-learning results, was computed with the left-out test set of the testing sample.

### Ablation studies

We performed two ablation studies to evaluate the importance of the input interaction network and the epigenomic features in modeling transcript isoform expression levels.

In the first ablation study, we trained ANN, Random Forest, and Linear Regression models to model transcript isoform expression levels using only the epigenomic features without the interaction network. For the ANN, we used the ANN function in the PyTorch package (v1.8.1), and performed training and testing across all 358 target samples with the same set of transcript isoforms as defined above for GCN. To ensure the optimal hyperparmeters were chosen for each model, we performed hyperparameter search using the independent validation set of transcript isoforms. Several hyperparameters were optimized by choosing from multiple values, including 1) dropout: 0.1, 0.2, 0.3 and 0.4, 2) learning rate: 1e-2, 1e-3, 1e-4 and 1e-5, 3) weight decay: 1e-3, 1e-4, 1e-5 and 1e-6, and 4) batch size: 16, 32, 64 and 128. We used 6 hidden layers each with 128 hidden nodes to closely mimic the number of parameters of our GCN models, and applied ReLU at each layer as the non-linear activation function. Similarly, we used the Adam optimizer and MSE as the loss function. Both GCN and ANN models had 118,145 trainable parameters. For the Random Forest and Linear Regression models, We used the RandomForestRegressor and LinearRegression functions in the Scikit-learn package (v1.0.1), and a setting of n estimators=100 for all our Random Forest models.

In the second ablation study, we replaced all epigenomic features with random Gaussian features such that the resulting GCN models were supplied only with the real interaction network as input but no real epigenomic features. The random features were generated as a tensor in the shape of (number of transcript isoform nodes, 147), where values were sampled from a standard Gaussian distribution with a mean of 0 and standard deviation of 1 using the randn function in the PyTorch package (v1.8.1).

### Meta-learning

We used a meta-learning approach to implement the LOSO, LOTO, and ST learning strategies. To model transcript isoform expression levels in sample *i*, we first formed the set of training samples *S* according to the learning strategy as explained in Results. Then for each transcript isoform *t*, we took its inferred expression level in sample *i* based on the model trained on each sample *j* in *S*. This inferred expression level is used as the “feature” of transcript isoform *t* in sample *i* contributed by sample *j*. For each sample not in *S*, this “feature” value was set to 0.

Secondary Linear Regression models were then trained to relate these “features” with the actual transcript isoform expression levels in sample *i*, using only transcript isoforms on the training chromosomes. We evaluated three different regularization methods, using the Lasso, Ridge, and ElasticNet functions in the Scikit-learn package (v1.0.1), with a setting of max iter=1000. For the linear regression model of each sample *i*, the value of the alpha parameter was determined using a grid search (with values 0.1, 0.01, 0.001, 0.0001, 0.00001, 1, 10, 100, 1000 and 10000), and the value that maximized the modeling performance of the transcript isoforms on the validation chromosomes was used.

Each of the trained models was applied to infer the expression levels of transcript isoforms on the testing chromosomes in sample *i* and compared with the actual measured levels to quantify its performance. For the three linear regression models (based on LASSO, Ridge regression, and Elastic Net), we selected the one with the best validation set performance and reported its test set performance.

### Feature importance scores

We used two different methods to evaluate the importance of each feature in a GCN model trained on data from a single sample, namely saliency attribution and feature value permutation. To explain these methods, we define the following notations. Let **X** *∈* R*^N×F^* be the node-feature matrix of the *N* nodes in the input graph of the GCN each with *F* features. Let *G_θ_*(**X**) *∈* R*^N^* be the output of these *N* nodes inferred by the GCN model *G_θ_* with trainable parameters *θ*. Finally, let *M ⊂ {*1*,…,N}* be the subset of nodes that correspond to the transcripts.

The saliency attribution method evaluates the gradient of the GCN output of a node with respect to a feature of it. Specifically, we applied Captum’s Saliency method (captum package v0.7.0) using a single backward pass:

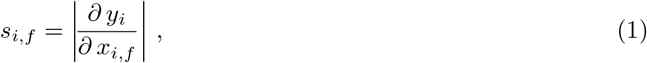

where *y_i_*is the GCN output of node *i* and *x_i,f_* is feature *f* of node *i*. The overall saliency of a feature in this GCN is then defined as the average across all transcript nodes:

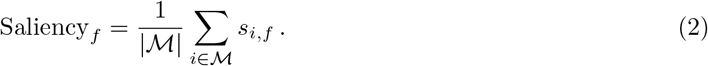

Finally, to compute a single saliency value for each feature across multiple GCN models, we performed log-10 transformation of the saliency value from each GCN, min-max normalized the log-transformed values per model to the [0,1] range, and then averaged across samples. A feature with a larger final value is considered more important.

The feature value permutation method evaluates the change of the loss when the values of a feature are permuted. For the real node-feature matrix **X**, the mean-squared error is defined as:

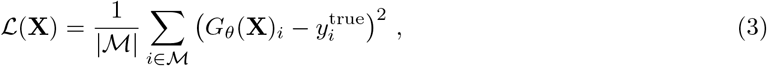

where *G_θ_*(**X**)*_i_* and *y*^true^ are the GCN-inferred and actual experimentally measured expression levels of transcript isoform *i*, respectively.

Now, suppose the values of feature *f* are randomly permuted to form the new permuted node-feature matrix **X**^(*f*)^. The permutation importance of feature *f* in this GCN is defined as the change of the loss due to this permutation:

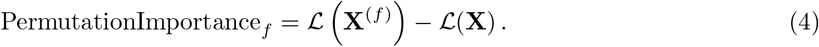

Finally, to compute a single permutation importance value for each feature across multiple GCN models based on their individual permutation importance values, we first added each of these permutation importance values by one, performed log-10 transformation of the absolute value of each of them and added back its sign, divided each resulting value by the maximum absolute value of them, and finally averaged these values. A feature with a larger final value is considered more important.

### Gene-level expression modeling and benchmarking

To benchmark our method against GC-MERGE ^21^ and Chromoformer ^9^ at the gene level, we converted all our transcript isoform-level predictions to a common gene-based representation and evaluated Pearson correlation on log2-transformed gene TPM values. For our method, transcript-level predictions were aggregated to gene TPM using length-weighted pseudo-counts derived from transcript logits, followed by renormalization to a total of 10^6^ TPM. For GC-MERGE and Chromoformer, we used their regression model outputs for each cell line and evaluated them under the same 7-fold chromosome-stratified crossvalidation framework used for our method, so that training and testing splits were matched across all three methods. This ensured that all comparisons were made on the same fold structure. To assess whether sex chromosomes altered the conclusions, we repeated the analysis on two gene sets: 7-fold cross validation test set with all annotated genes, and the same test set excluding chrX and chrY. In both cases, Pearson correlation was computed after applying log_2_(TPM + 10*^−^*^3^) to the true and predicted values.

### Analysis of challenging and non-challenging genes

We performed a stratified analysis of model performance under a z-score expression threshold. For each sample, transcript expression labels were standardized using a z-score computed across all transcript labels in that sample. Transcripts with *z ≥* 1.5 or *z ≥* 2.0 were considered decently expressed in two respective settings. For each gene-sample pair, the number of decently expressed isoforms was counted, and pairs with more than three such isoforms were classified as challenging, whereas pairs with three or fewer were classified as non-challenging. For every gene-sample pair, we computed Pearson correlation between the predicted logits and the measured transcript labels. Gene-sample pairs were then stratified by the gene’s total annotated transcript count. Within each transcript-count stratum, we calculated the mean Pearson correlation separately for challenging and non-challenging pairs.

### Investigating epigenetic feature divergence among transcript isoforms of the same gene

To assess whether transcript isoforms from the same gene are effectively redundant in epigenetic feature space, we constructed pairwise difference vectors from the full transcript-level epigenetic representation. The feature space comprised 21 genomic positions per assay (Up5-Up1, TSS1-TSS5, first exon/intron, internal exons/introns, last intron/exon, and Dw1-Dw5) across seven epigenetic tracks: H3K27ac, H3K27me3, H3K36me3, H3K4me1, H3K4me3, H3K9me3, and DNA methylation. Transcripts with any missing epigenetic feature encoded as-1 were excluded. For each isoform pair, we computed the absolute difference vector across all features and standardized each feature before PCA (z-scoring across pairs). We then projected these pairwise difference vectors onto the first two principal components and compared same-gene with cross-gene pairs in setting 1, and high-versus-low expression-divergence pairs within genes versus the corresponding cross-gene comparison in setting 2. In the expression divergence analysis, for each sample, transcript isoform pairs were identified within the same gene, and the withinsample expression difference between the two isoforms was calculated from their corresponding expression values. The paired epigenetic feature vectors were obtained from the same sample, and epigenetic divergence was quantified as the Euclidean distance between the 21 epigenetic features across all epigenetic marks. Isoform-pair observations were subsequently stratified into low-and high-expression-difference groups based on the median within-sample expression difference, and the distributions of epigenetic divergence were compared between groups. In addition, cosine similarity between paired epigenetic feature vectors was calculated to assess the degree of feature redundancy among isoforms. Feature-specific analyses further quantified Euclidean distances separately across individual epigenetic marks and genomic positions. Differences between groups were evaluated using two-sided Mann–Whitney U tests.

## Acknowledgments

KYY is supported by National Cancer Institute of the National Institutes of Health under Award Numbers P30CA030199, R01CA287114, and R01CA296261, National Heart, Lung, and Blood Institute of the National Institutes of Health under Award Number R21HL177724, National Institute on Aging of the National Institutes of Health under Award Numbers R01AG085498 and U54AG079758, National Institute of General Medical Sciences of the National Institutes of Health under Award Number R21GM159319, V Foundation for Cancer Research under Award Number V2025-028, and internal grants of Sanford Burnham Prebys Medical Discovery Institute. The content is solely the responsibility of the authors and does not necessarily represent the official views of the National Institutes of Health. SHC was supported by the California Institute for Regenerative Medicine under a pre-doctoral fellowship (EDUC4-12813).

## Contributions

KYY conceived the study. SHC and CHS processed the data. SHC implemented the methods and performed the computational modeling. SHC, CHS, QC, AD, and KYY performed the data analysis and interpreted the results. SHC and KYY wrote the manuscript. All authors read and approved the final version of the manuscript.

## Supplementary information

### Supplementary Text

#### Investigation of additional factors that affect modeling results

To better understand why the inferred transcript isoform expression levels were less accurate when applying a model to another sample, we examined a variety of potential factors. First, we found that transcript isoform expression levels in immortalized cell lines were inferred more accurately, regardless of the type of the training sample (Figure S5a). Similarly, transcript isoform expression levels in cell samples were inferred more accurately than those in tissue samples, regardless of whether a cell or tissue sample was used in training the model (Figure S5b). Third, although all transcript isoform expression levels were obtained by RNA-seq, two different RNA-seq protocols were used, namely mRNA-seq that uses poly-A enrichment to focus on protein-coding transcripts, and total RNA-seq that provides a more uniform coverage of different transcript types. The modeling results show that transcript isoform expression levels measured using total RNA-seq were inferred more accurately, no matter expression data of the training sample were obtained using mRNA-seq or total RNA-seq (Figure S5c). Fourth, WGBS was also performed using two different protocols, namely a more standard protocol and an amplification-free protocol, Post-Bisulfite Adaptor Tagging (PBAT) ^40^. We found that when DNA methylation in the testing sample was measured using PBAT, the inferred transcript isoform expression levels were more accurate, regardless of the WGBS protocol used to measure DNA methylation in the training sample (Figure S5d). Fifth, we found that transcript isoform expression levels of males were more accurately inferred regardless of the sex of the training sample (Figure S5e). Sixth, we found that transcript isoform expression levels of non-disease samples were more accurately inferred regardless of the disease status of the training sample (Figure S5f). Finally, some “samples” in our core dataset were actually multiple samples pooled together. We found that when both the training and testing samples were such pooled samples, the modeling performance was higher than the other cases (Figure S5g).

We next performed a regression analysis that uses all these factors of the training and testing samples together as the explanatory variables and the resulting modeling performance as the response variable. The results (Figure S6) show that the biomaterial type of the testing sample has the largest effect size on modeling performance, followed by the disease status of the testing sample, while all the other variables have smaller effect sizes.

## Supplementary Figures

**Supplementary Figure S1:**
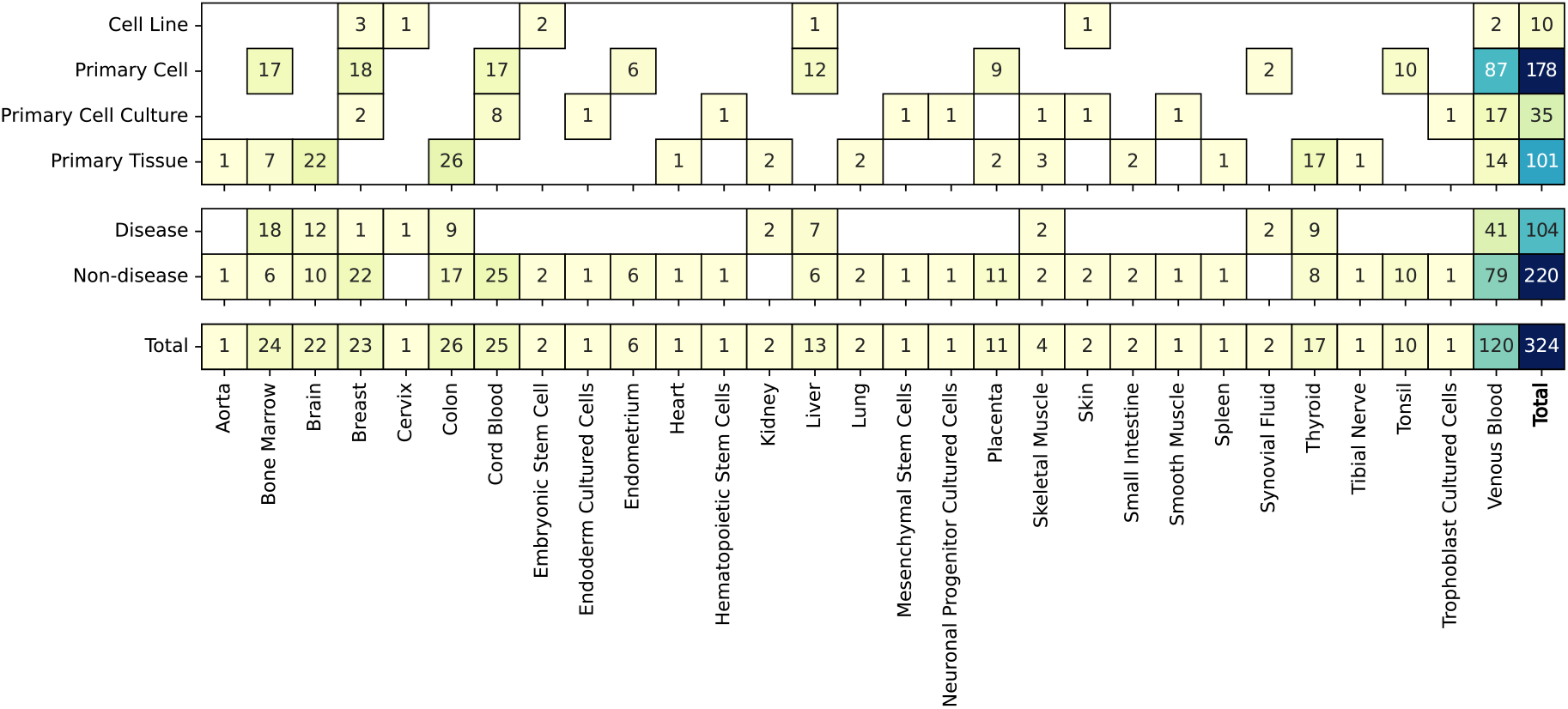
Summary of the samples in our core dataset. The samples are grouped by the tissue type (columns), biomaterial type (first block of rows), and disease state (second block of rows). Each table entry shows the number of samples in each group.

**Supplementary Figure S2:**
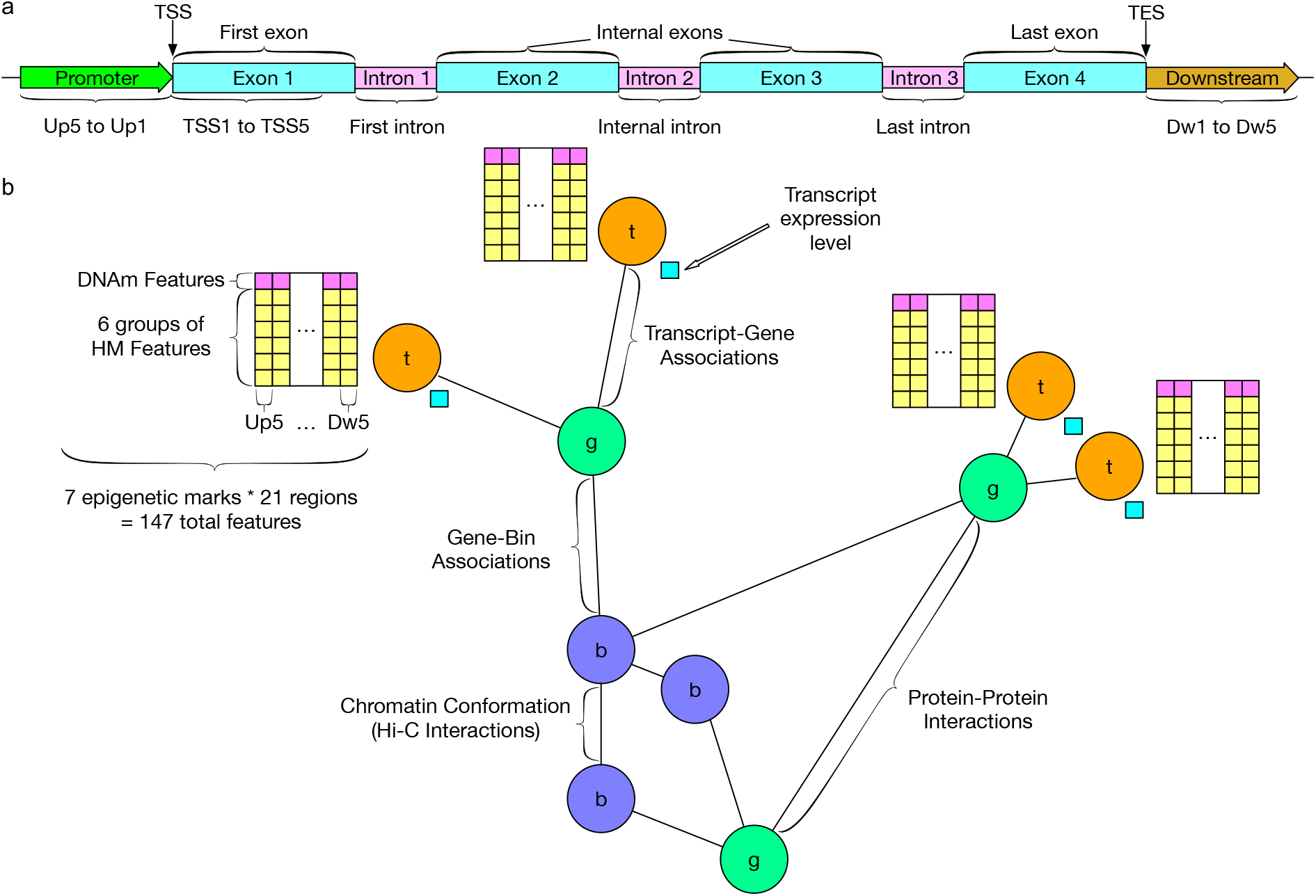
Data representation for modeling transcript isoform expression levels. a, The 21 regions defined according to the structure of each transcript isoform, based on which epigenomic features are computed. The 21 regions include 5 upstream regions (Up1-Up5), 5 regions immediately after the transcription start site (TSS1-TSS5), 6 gene body regions (first exon, first intron, internal exons, internal introns, last intron, and last exon), and 5 downstream regions (Down1-Down5). **b,** The integrated interaction network that links genes and their protein products (both denoted as “g”), transcript isoforms (“t”), and genomic bins (“b”).

**Supplementary Figure S3:**
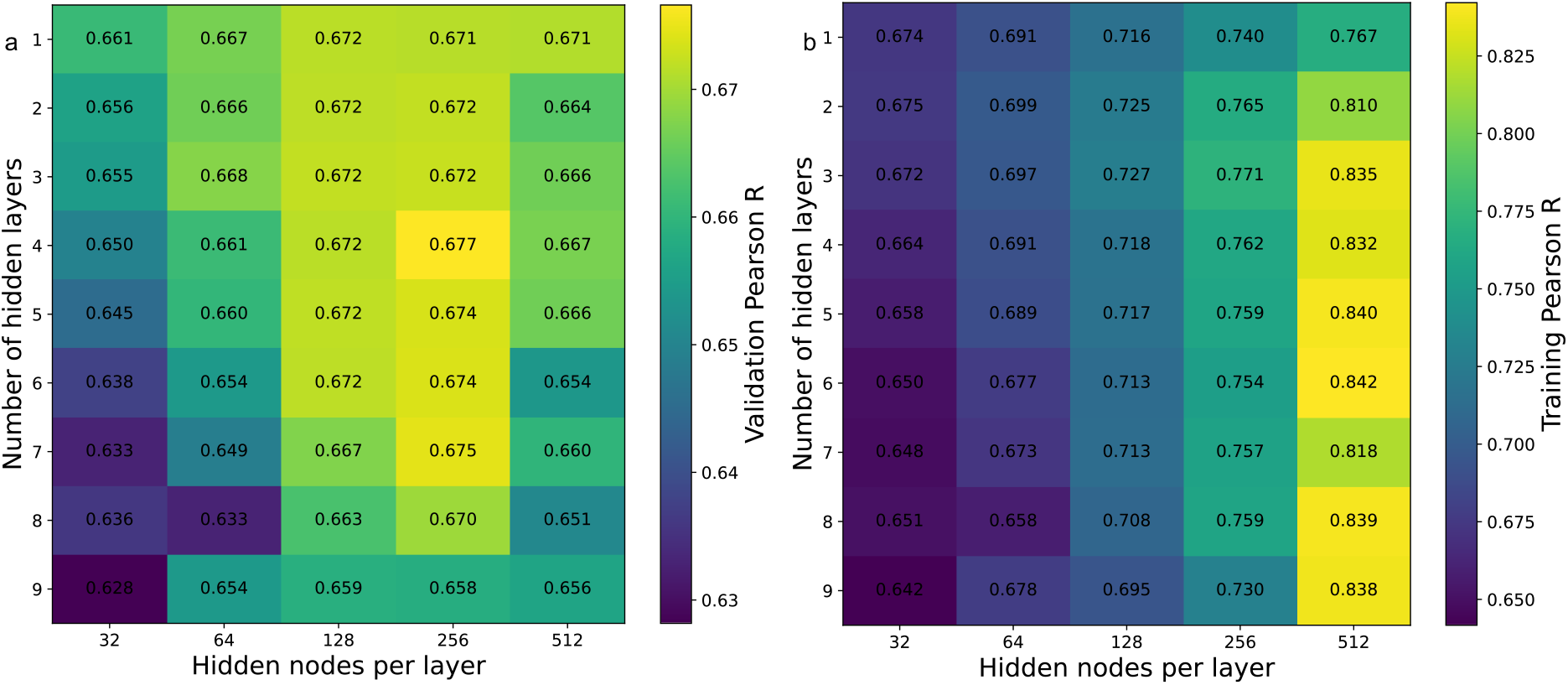
Hyperparameter search for GCN architecture. a-b, Pearson correlation performance of GCN models with different hidden-layer width (32, 64, 128, 256 and 512 nodes) and depth (1 to 9 layers), quantified by validation (a), and training (b) set performance.

**Supplementary Figure S4:**
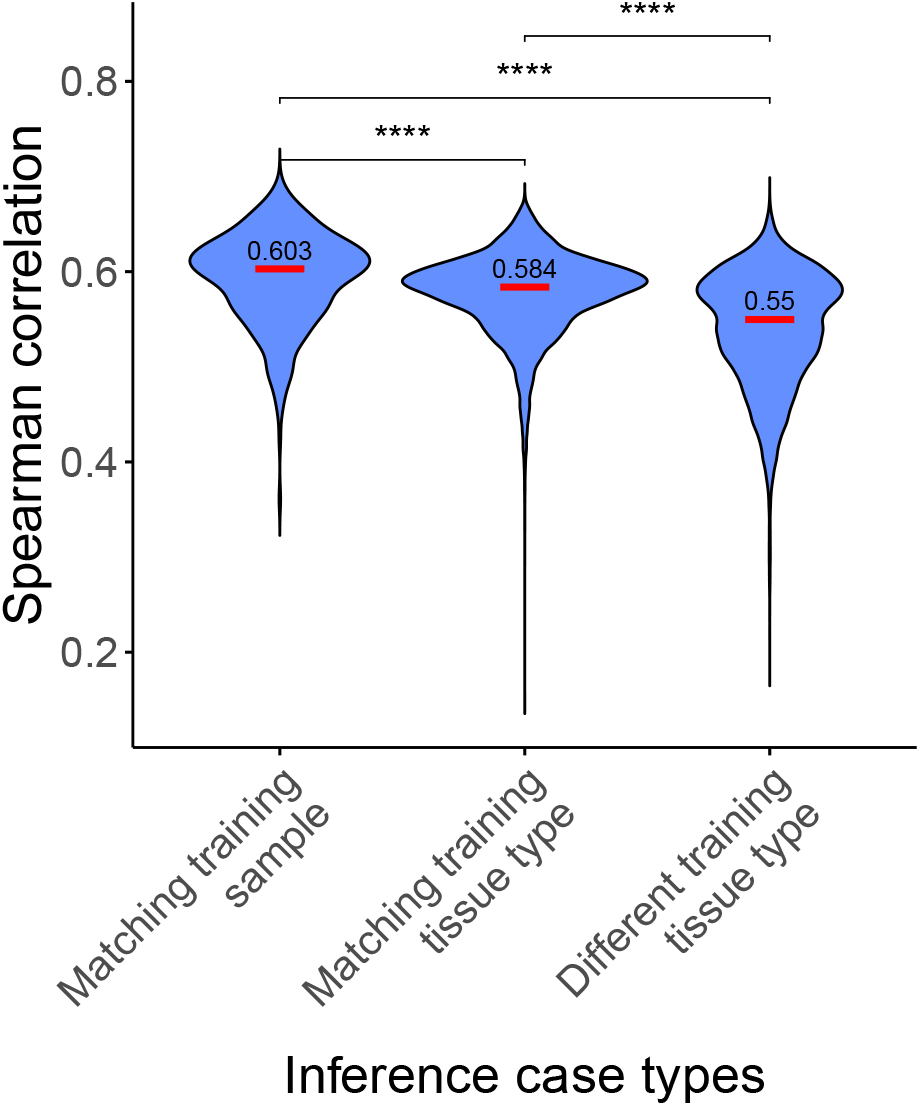
Modeling performance using training data from one sample at a time, quantified by an alternative performance measure. Distribution of modeling performance (quantified by Spearman correlation) when the training and testing samples were either the same, different but from the same tissue type, or from different tissue types. P-values were calculated using two-sided Wilcoxon rank-sum test. *:p*<*0.05; **:p*<*0.01; ***:p*<*0.001; ****:p*<*0.0001.

**Supplementary Figure S5:**
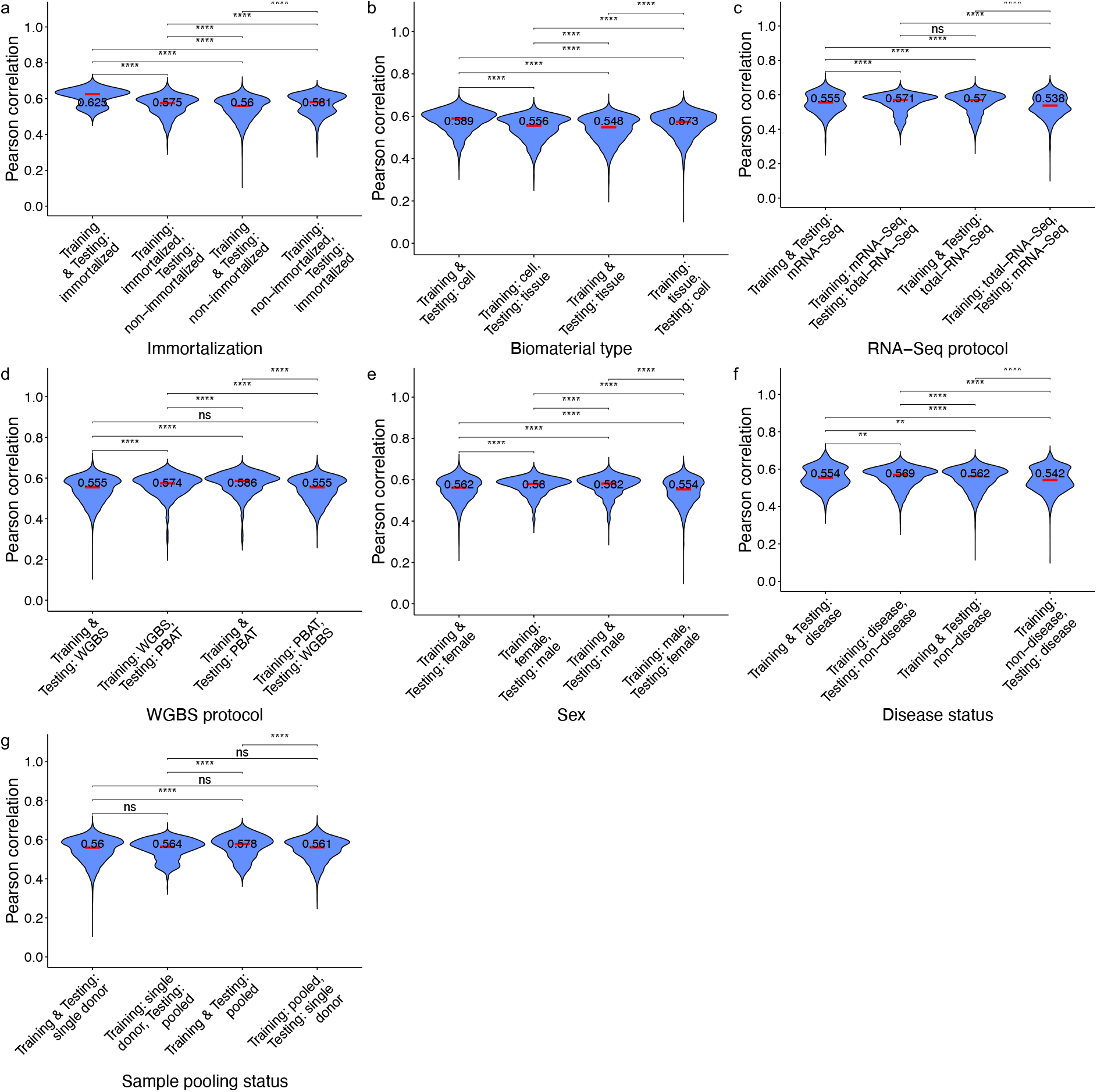
Modeling performance using training data from one sample at a time, stratified by additional factors. a-g, Distribution of modeling performance when the training and testing samples have matched or unmatched immortalization status (a), biomaterial type (b), RNA-seq protocol (c), WGBS protocol (d), sex (e), disease status (f), or sample pooling status (g). P-values were calculated using two-sided Wilcoxon rank-sum test. *:p*<*0.05; **:p*<*0.01; ***:p*<*0.001; ****:p*<*0.0001; ns:not significant.

**Supplementary Figure S6:**
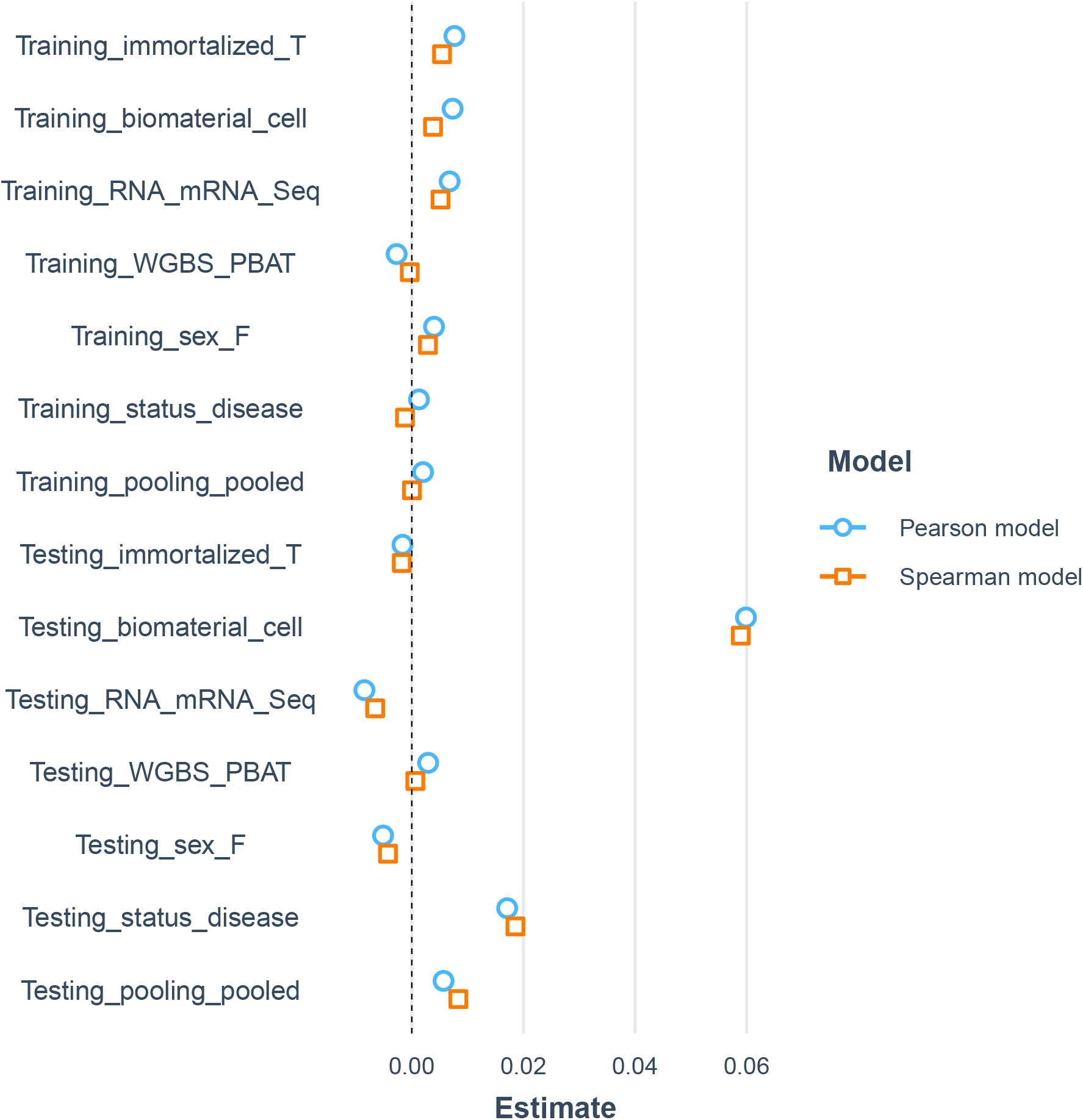
Joint effect of additional factors on modeling performing when using training data from one sample at a time. The X-axis shows the estimated effect size of each binary variable in the multivariate regression model that explains modeling performance, quantified by Pearson correlation or Spearman correlation. The label of each variable consists of three parts, namely 1) whether the variable is about the training or testing sample, 2) the factor of concern, and 3) the value of the factor that defines the positive direction. For example, “Training immortalized T” denotes the binary variable that the training sample is immortalized cells (where “T” means true), and a positive effect size of this variable means that when the training sample is immortalized cells, the modeling performance tends to be higher than when this is not true.

**Supplementary Figure S7:**
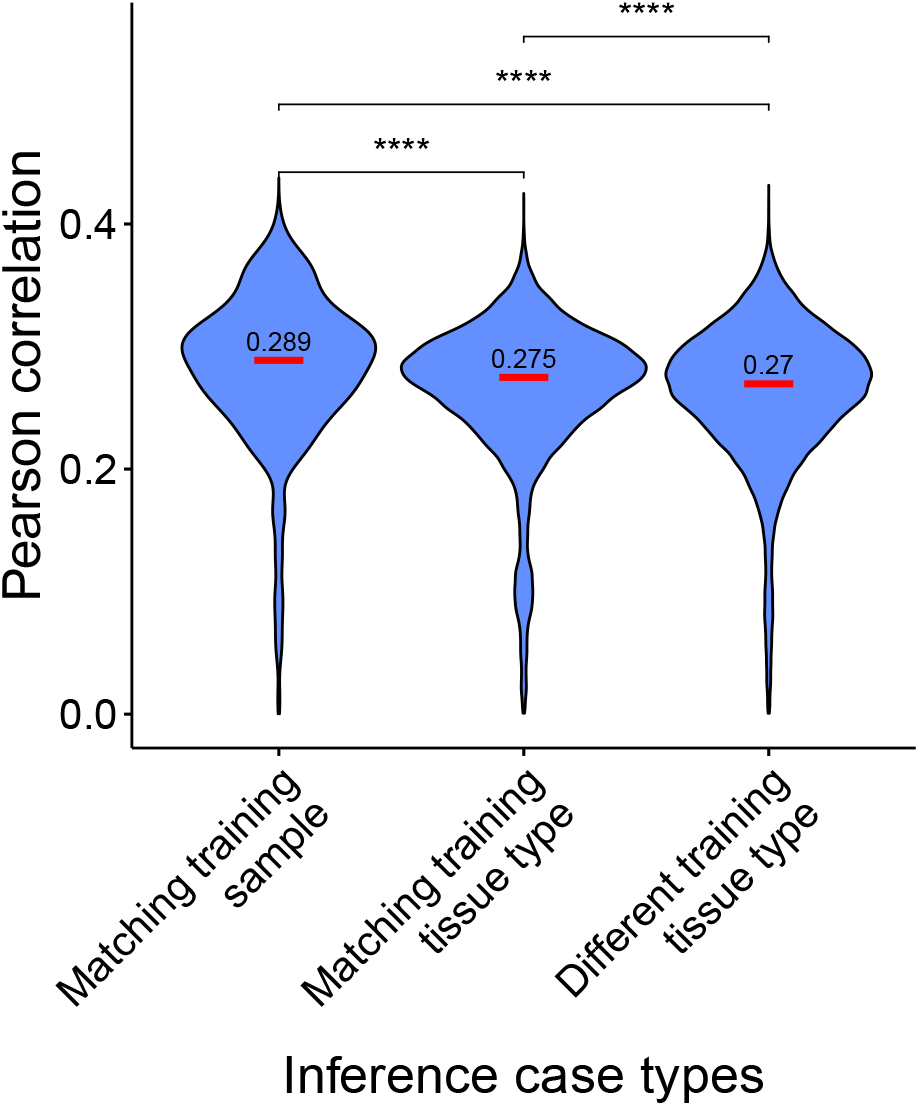
Modeling performance using training data from one sample at a time, with intentional random epigenomic features. Distribution of modeling performance when the training and testing samples were either the same, different but from the same tissue type, or from different tissue types, with intentional random epigenomic features. P-values were calculated using two-sided Wilcoxon rank-sum test. *:p*<*0.05; **:p*<*0.01; ***:p*<*0.001; ****:p*<*0.0001.

**Supplementary Figure S8:**
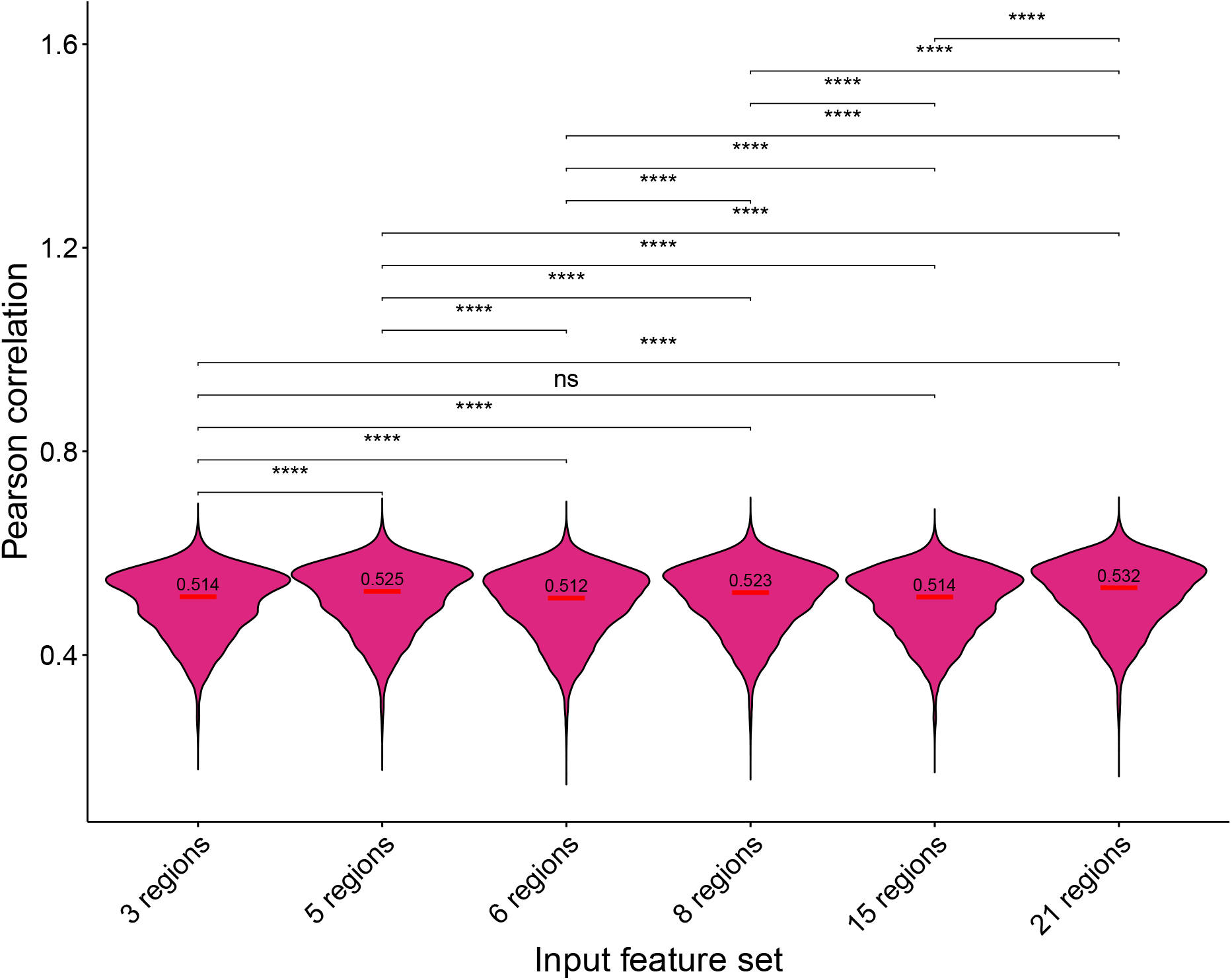
Comparing models with different input feature set granularity. Performance of RF models for our original feature definitions with 21 regions and those with 3 regions (promoter, exons, and introns), 5 regions (promoter, exons, introns, whole gene body, and downstream region), 6 regions (original gene body regions, namely first exon, first intron, internal exons, internal introns, last intron, and last exon), 8 regions (promoter region, original gene body regions, and down-stream region) and 15 regions model (original fix-sized regions, namely 5 upstream regions, 5 regions immediately after the transcription start site, and 5 downstream regions). P-values were calculated using two-sided Wilcoxon rank-sum test. *:p*<*0.05; **:p*<*0.01; ***:p*<*0.001; ****:p*<*0.0001; ns:not significant.

**Supplementary Figure S9:**
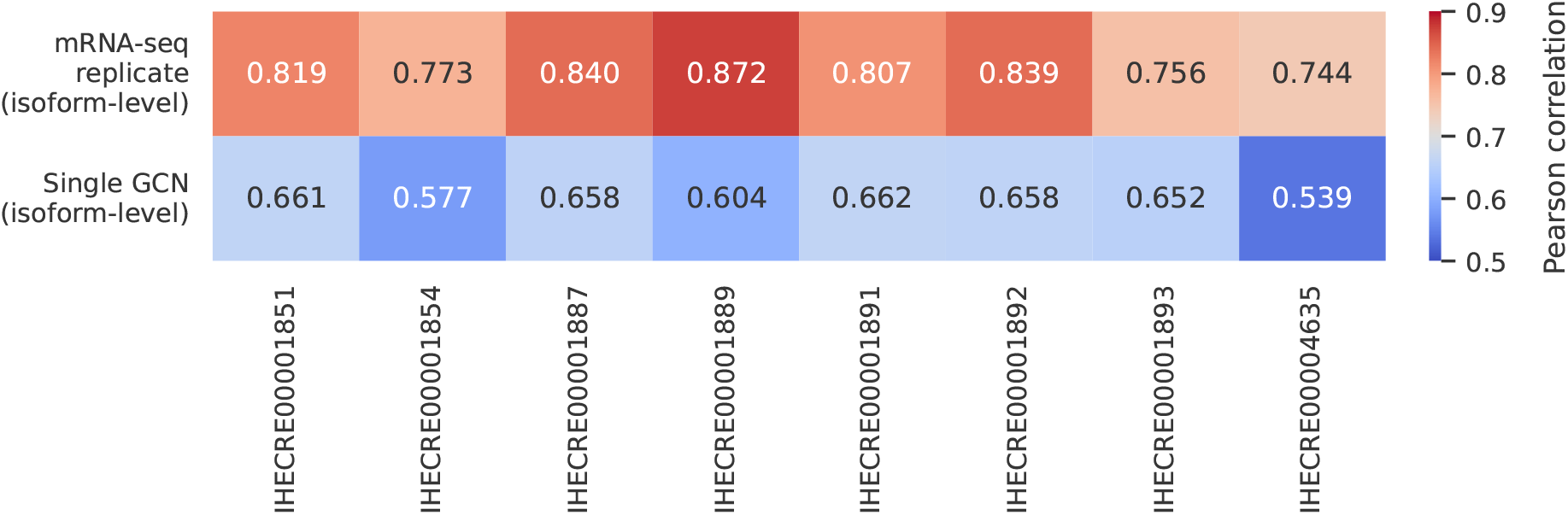
Assessing how close isoform-level modeling performance is from the maximum possible performance based on samples with replicated data produced using the mRNA-seq protocol. The two rows show the correlation of transcript isoform expression levels in the test set chromosomes between the two replicates (first row) or between the measured and GCN-inferred levels (second row).

**Supplementary Figure S10:**
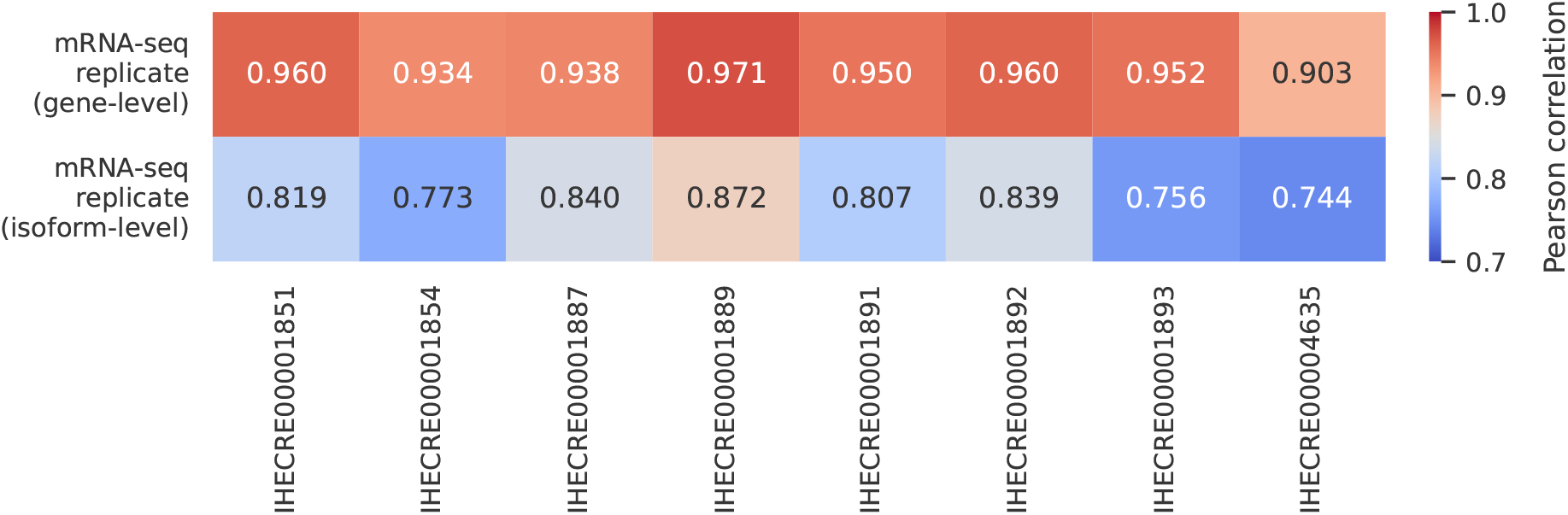
Comparing gene-level and isoform-level quantification concordance between replicates of mRNA-seq dataset from each sample. The first row shows the correlation of gene-level expression values in the testing-set chromosomes between the two replicates, and the second row shows the correlation of isoform-level replicates.

**Supplementary Figure S11:**
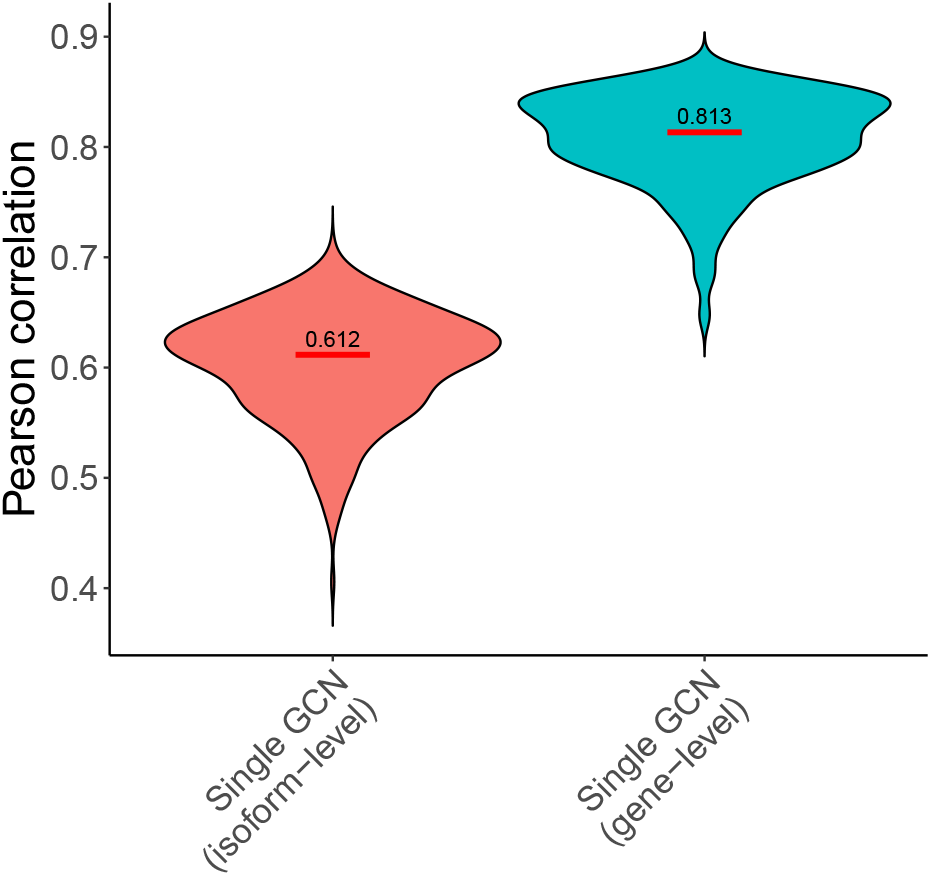
Gene-level performance of aggregated isoform-level modeling results. Distribution of isoform-level modeling performance compared to gene-level performance when the training and testing samples were the same. Pearson correlations were computed after aggregating transcript-level predictions to the gene level using isoform-length-weighted averaging.

**Supplementary Figure S12:**
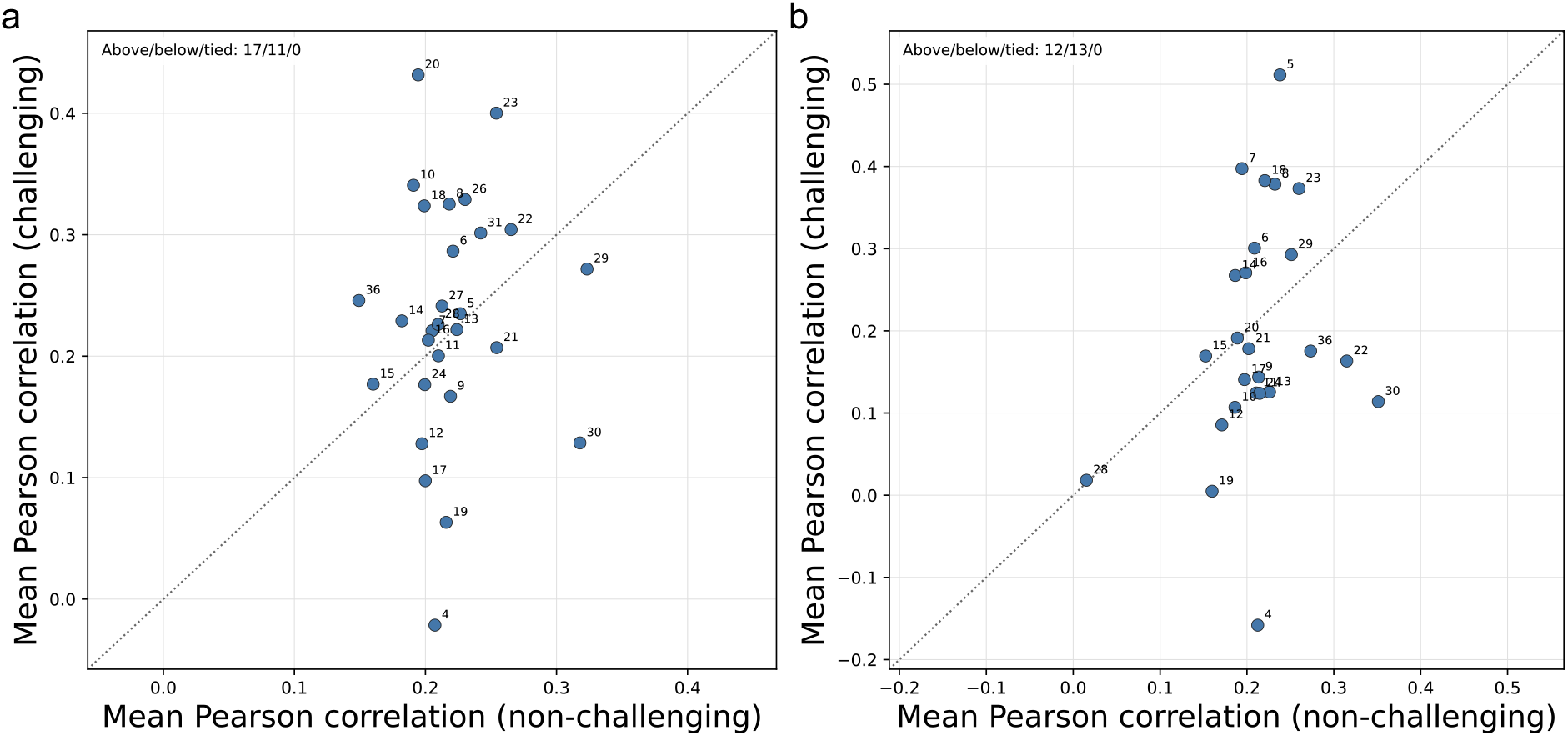
Modeling performance for challenging and non-challenging gene pairs under the isoform-complexity z-score criterion. a-b, Each dot denotes a gene group stratified by the number of transcript isoforms. The x-axis shows the mean Pearson correlation for non-challenging cases and the y-axis shows the mean Pearson correlation for challenging cases, with the threshold defined by *z* = 1.5 (a), and *z* = 2.0 (b).

**Supplementary Figure S13:**
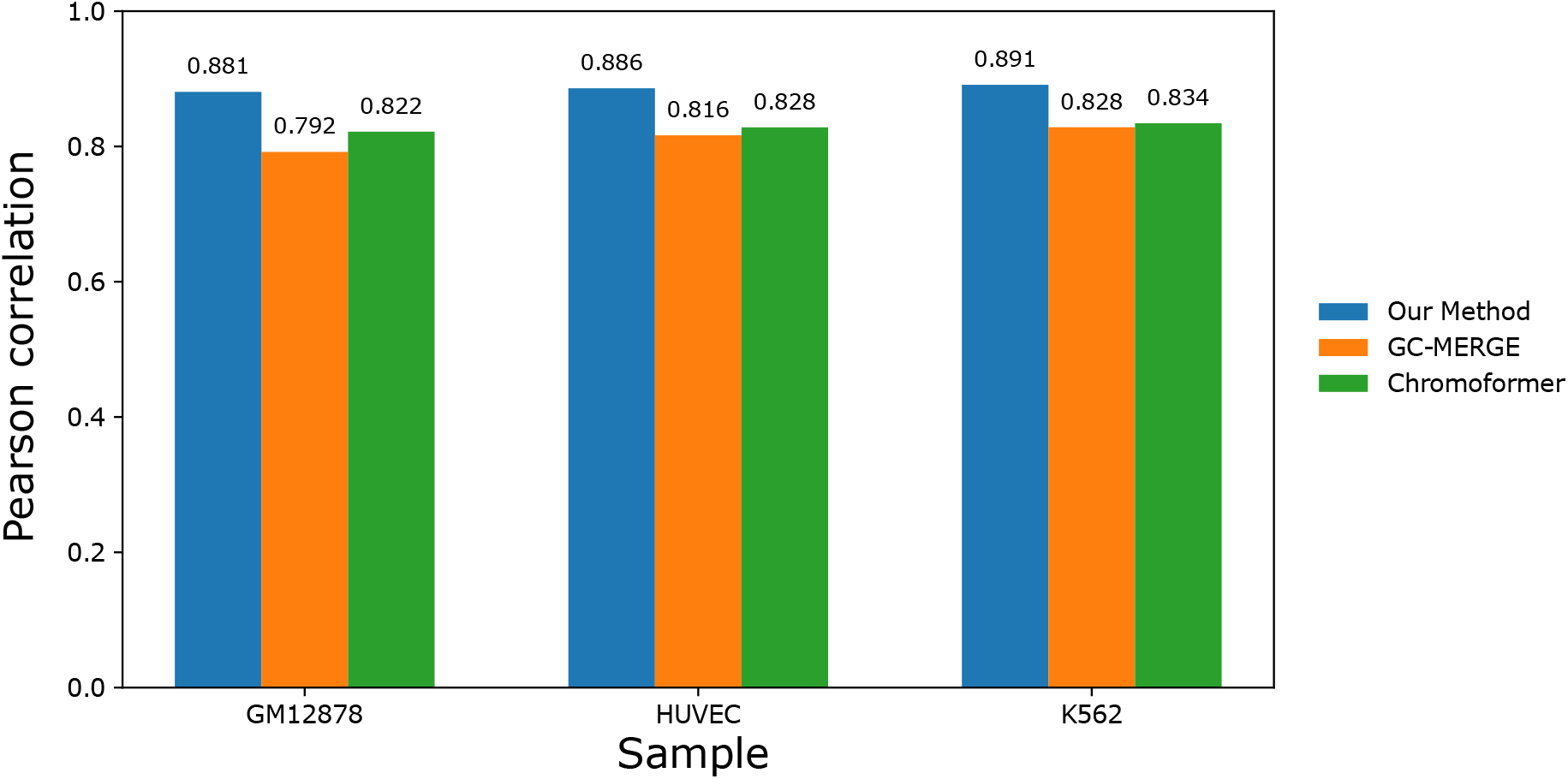
Gene-level modeling benchmark against GC-MERGE and Chromoformer. Pearson correlations are shown for the three cell lines under the same 7-fold chromosome-stratified evaluation. GC-MERGE and Chromoformer are GCN-based and transformer-based approaches performing gene-level modeling. We compared our gene-level values aggregated from isoform-level modeling and used a common set of genes across all 3 methods.

**Supplementary Figure S14:**
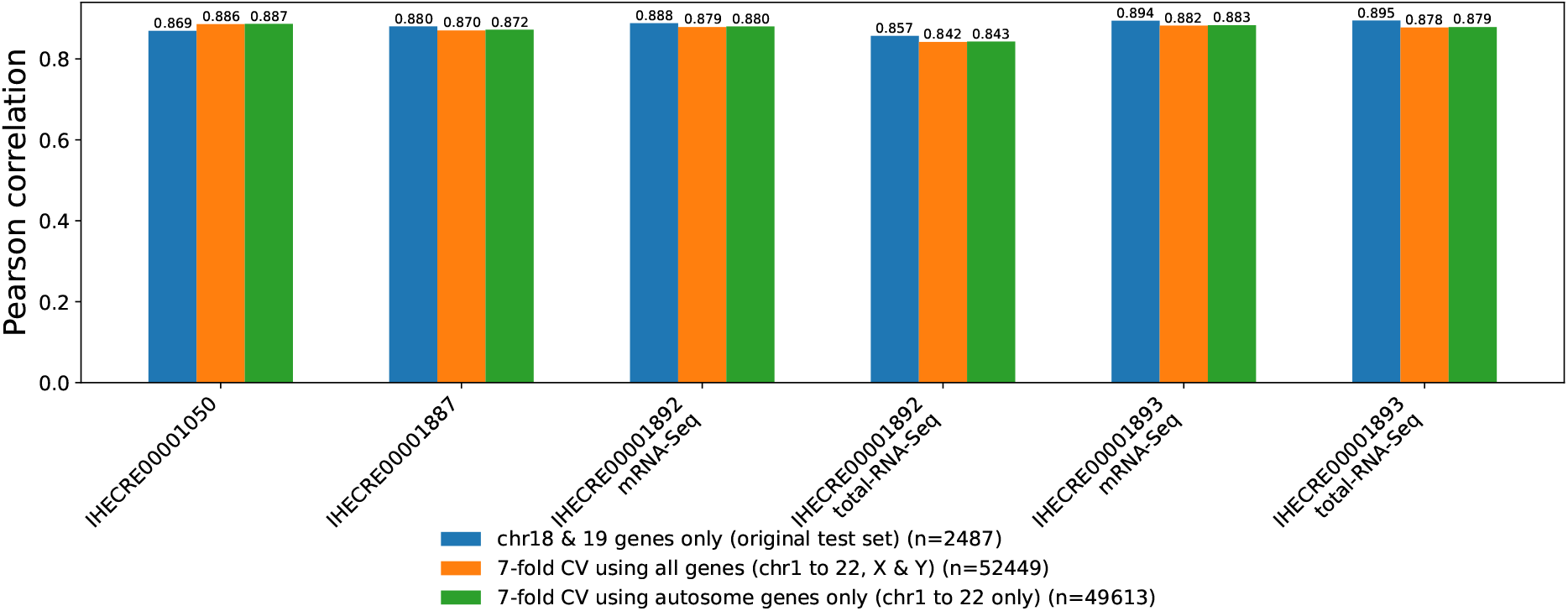
Testing whether sex chromosomes alter gene-level modeling results. Pearson correlations are shown for each sample when chrX and chrY are included or excluded from our 7-fold cross validation test set, as compared to the original test set (chr18 and chr19 genes only).

**Supplementary Figure S15:**
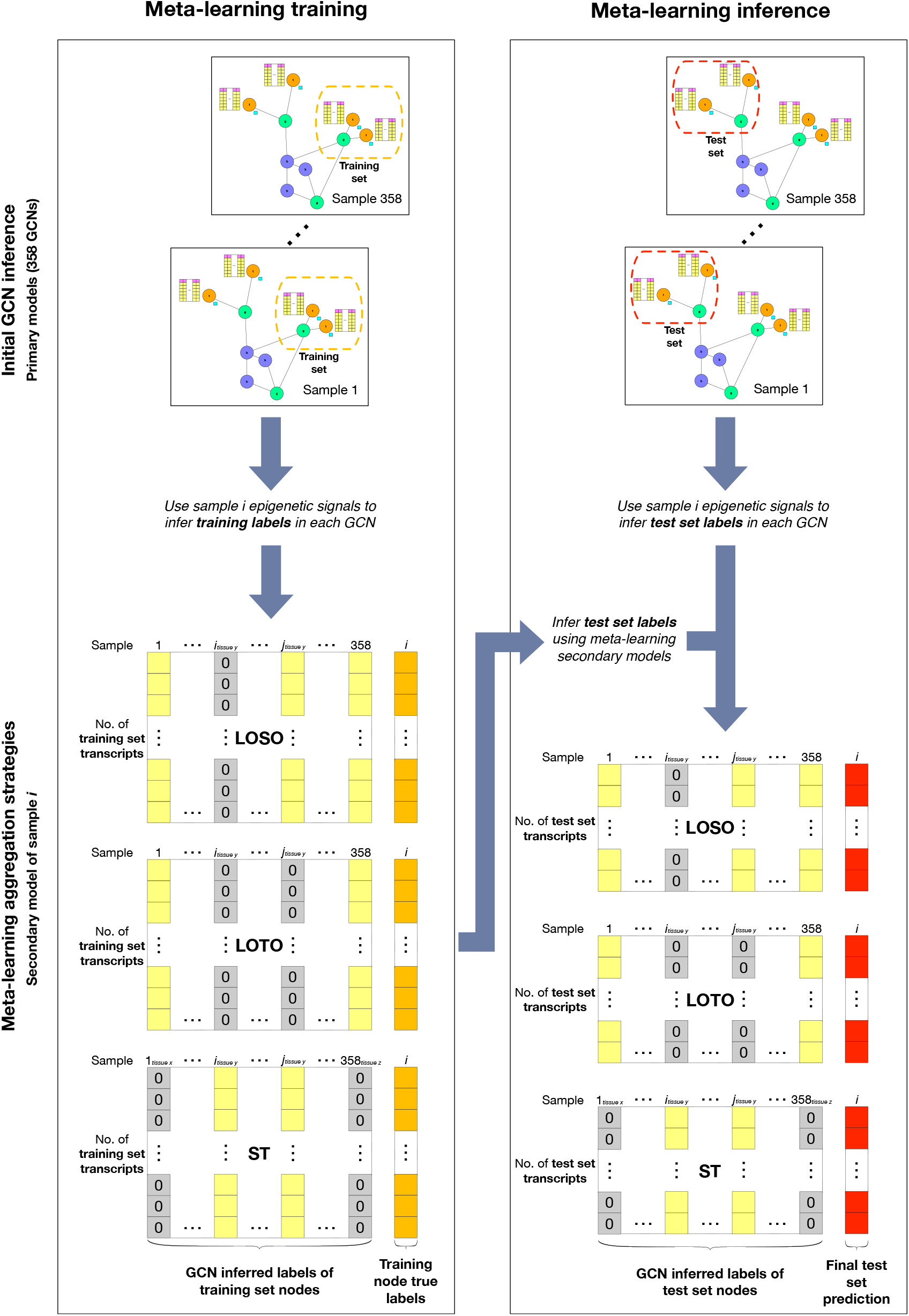
Schematic figure of the meta-learning strategy that combines information from multiple training samples to infer transcript isoform expression in a testing sample. To implement the three learning strategies, when a sample is not involved in meta-learning (i.e., it is not in the sample set *S*), it is replaced by a dummy predictor that infers all transcript isoforms in the target sample to have zero expression. As a result, in the LOSO, LOTO, and ST strategies, the samples replaced by the dummy predictor include the testing sample itself, all samples that share the same tissue type as the testing sample, or all samples that do not share the same tissue type as the testing sample, respectively.

**Supplementary Figure S16:**
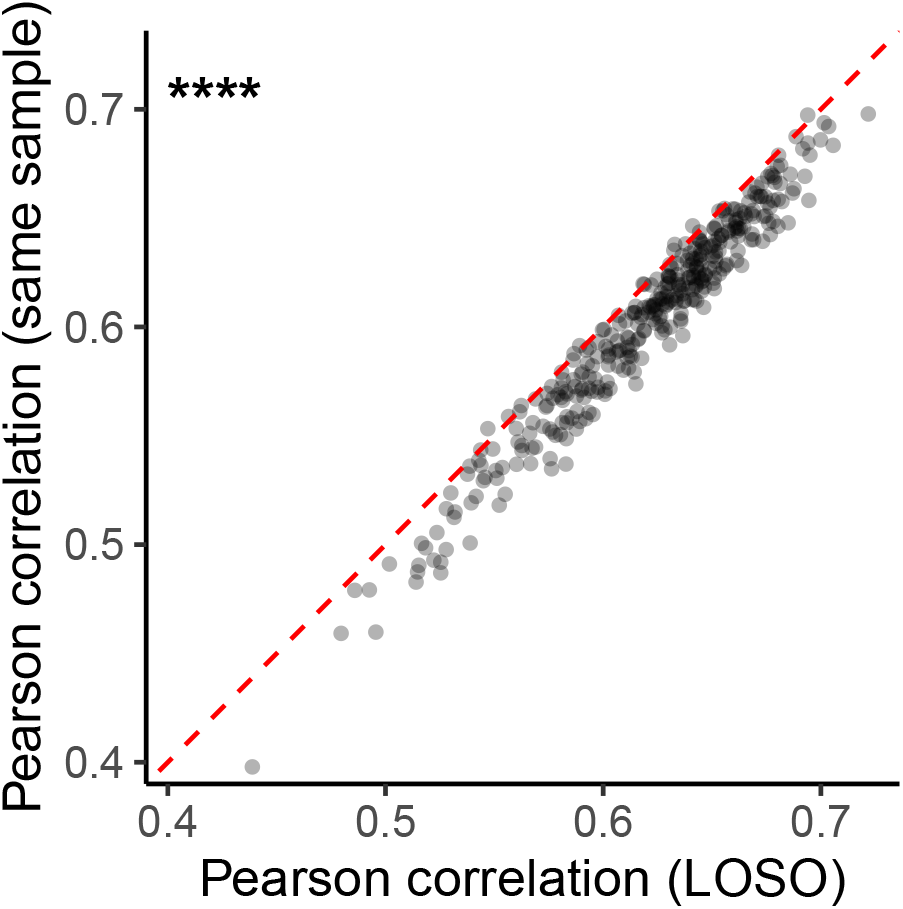
Comparing modeling performance of single-sample models trained on data from the testing sample with secondary models produced using the LOSO meta-learning strategy. P-value was calculated using two-sided Wilcoxon signed-rank test. *:p*<*0.05; **:p*<*0.01; ***:p*<*0.001; ****:p*<*0.0001.

**Supplementary Figure S17:**
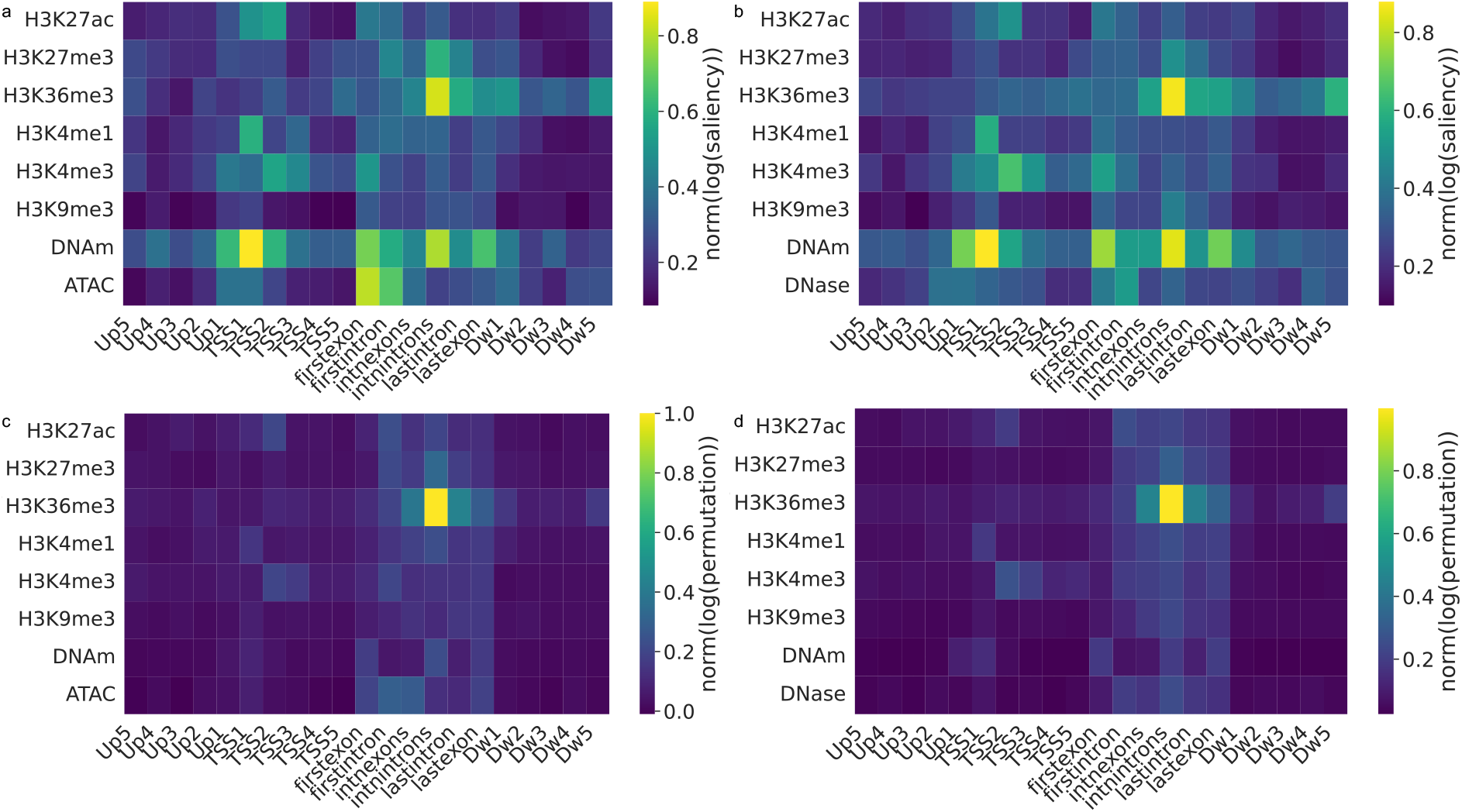
Effects of incorporating additional chromatin accessibility features into the single-sample models. a-d, Importance scores of features derived from the original seven types of epigenomic features and those derived from either ATAC-seq data only (a and c) or DNase-seq data only (b and d). Feature importance is quantified by saliency (a-b) or feature value permutation (c-d).

**Supplementary Figure S18:**
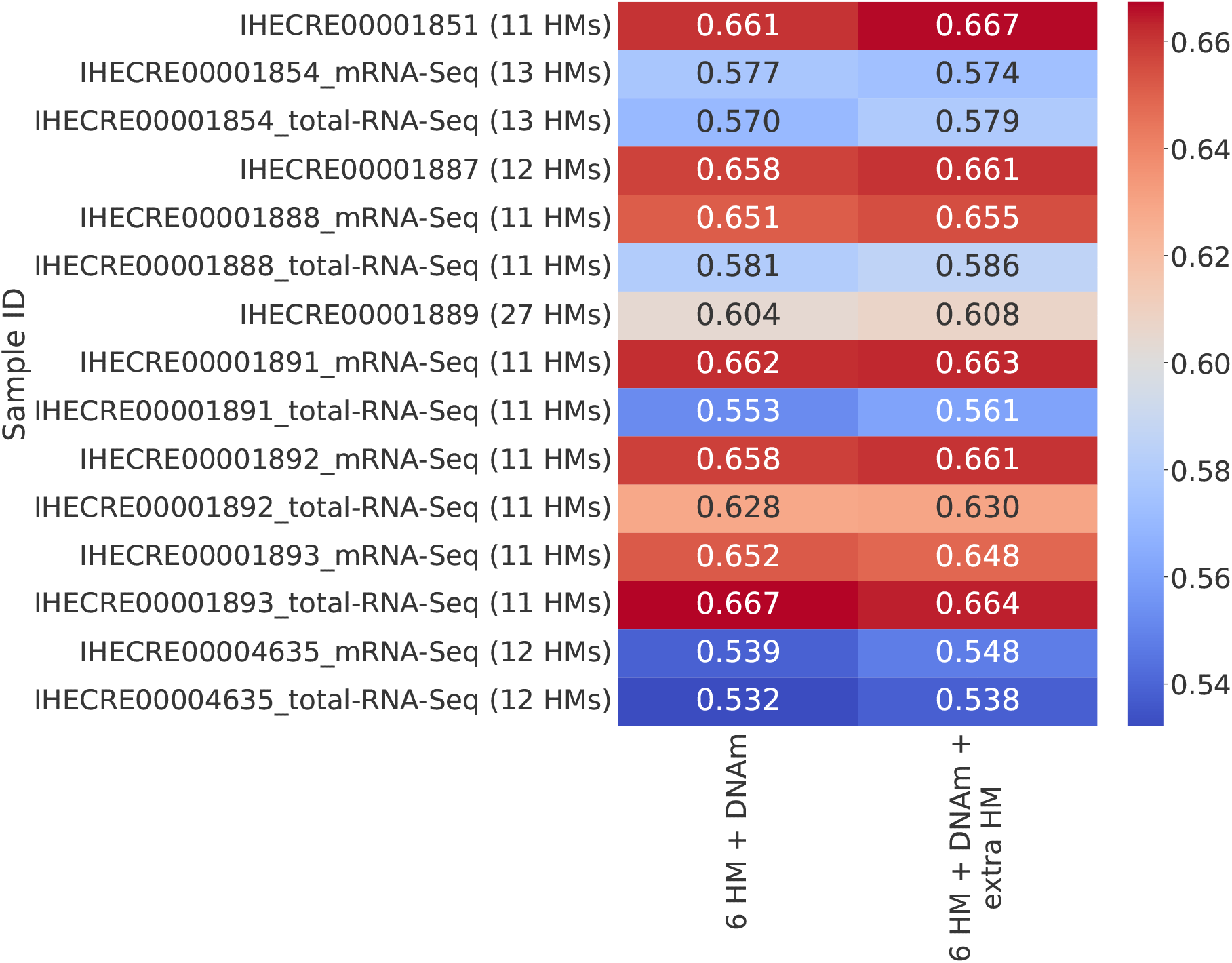
Comparing single-sample models trained with different number of histone modification datasets. Performance of models for our original feature definitions with 6 core histone marks and DNA methylation (“6 HM + DNAm”), or addition of extra histone modification datasets (“6 HM + DNAm + extra HM”). For the different samples, there were data for 5 to 21 additional types of histone modification, leading to the total of 11 to 27 types of histone modification as shown in parenthesis.

**Supplementary Figure S19:**
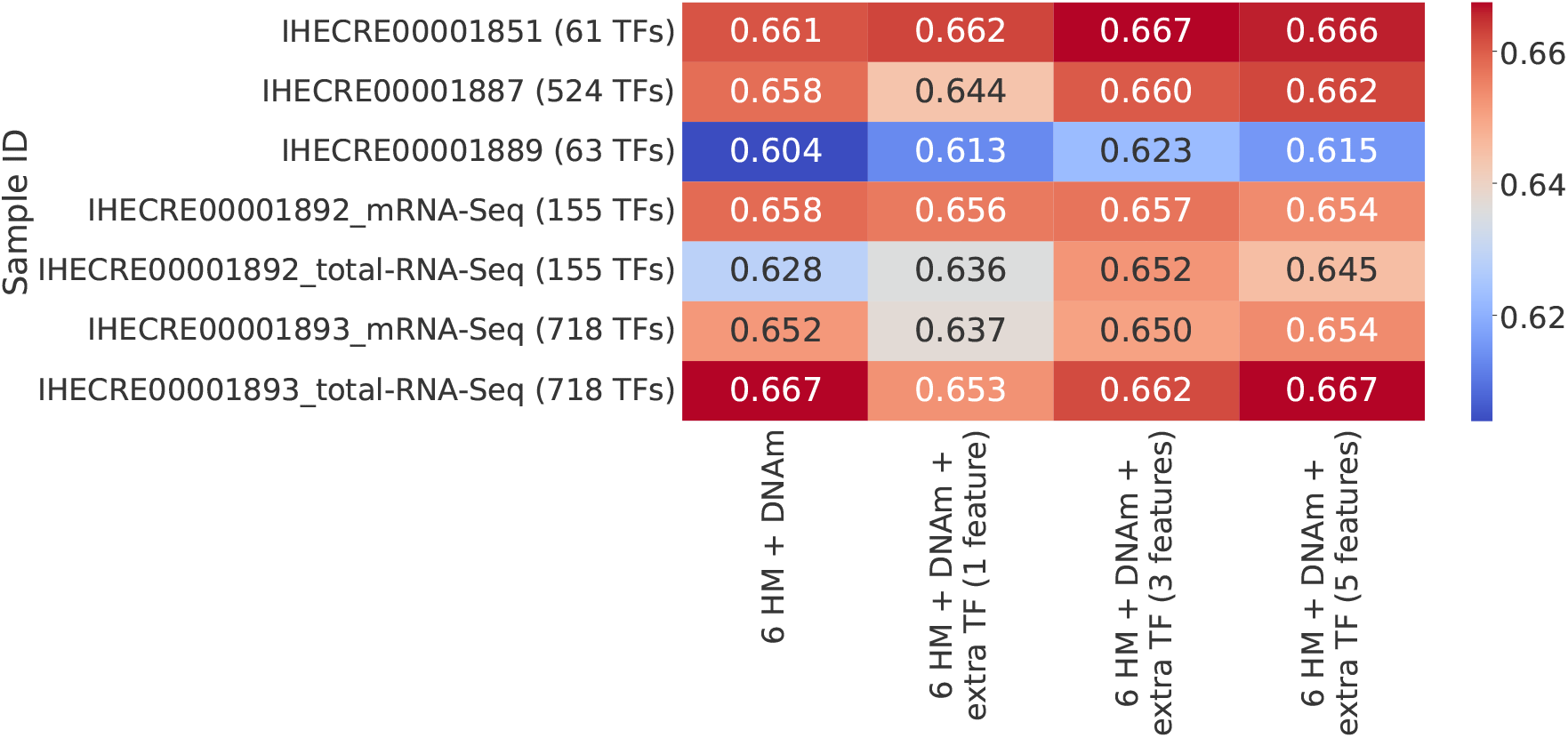
Comparing models trained with different number of transcription factor datasets. Performance of models for our original feature definitions with 6 core histone marks and DNA methylation (“6 HM + DNAm”), or addition of extra transcription factor (TF) datasets (“6 HM + DNAm + extra TF (x features)”, where x can be 1, 3, or 5). 1 feature: promoter only (1000 bp upstream and 500 bp downstream of TSS); 3 features: promoter, exon and intron; 5 features: promoter, exon, intron, full gene body and downstream (1000 bp downstream of TES).

**Supplementary Figure S20:**
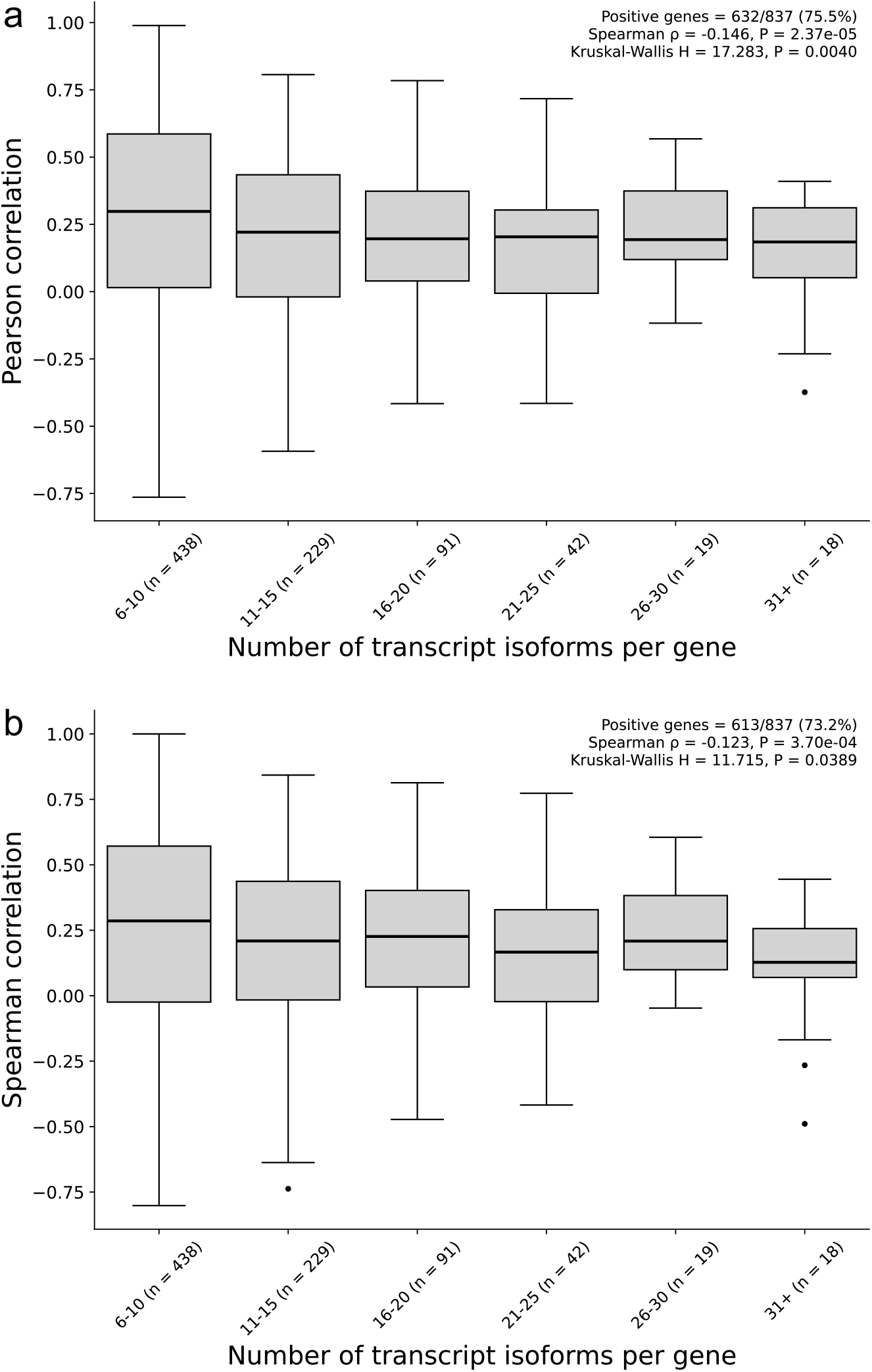
Within-gene isoform modeling remains positive after binning genes by isoform count. a, Pearson correlation versus the binned number of transcript isoforms. **b,** Spearman correlation versus the binned number of transcript isoforms. Each bin is annotated with the number of genes it contains, and the smallest bin is omitted from the figure due to high correlation instability when number of isoforms is low.

**Supplementary Figure S21:**
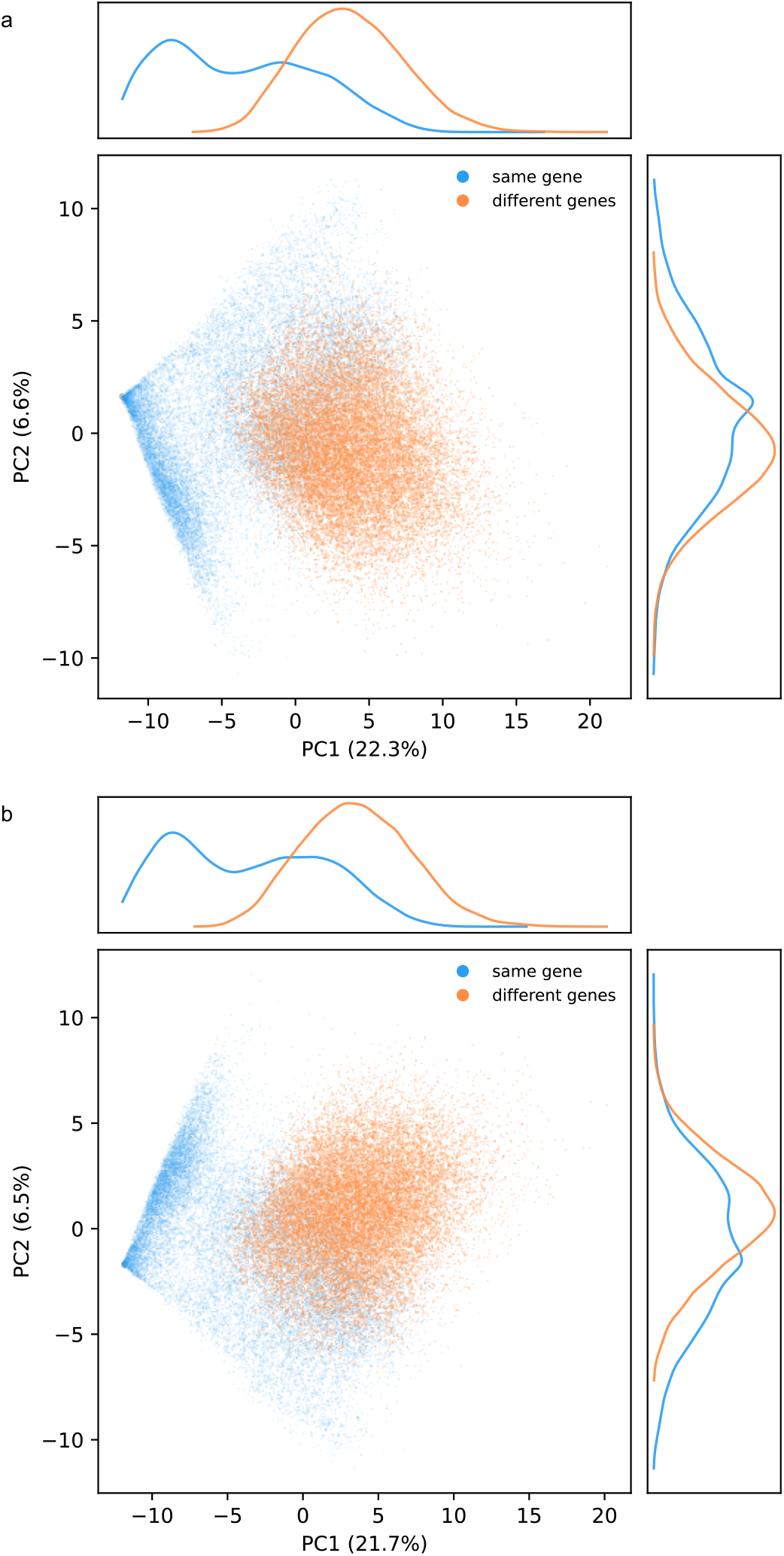
PCA of 147-feature epigenetic-divergence profiles for isoform pairs. a, Same-gene isoform pairs versus pairs drawn from different genes. **b,** Same-gene pairs stratified by expression divergence versus the corresponding cross-gene reference pairs. The 147-feature epigenetic-divergence profiles show considerable overlap, indicating substantial within-gene heterogeneity.

**Supplementary Figure S22:**
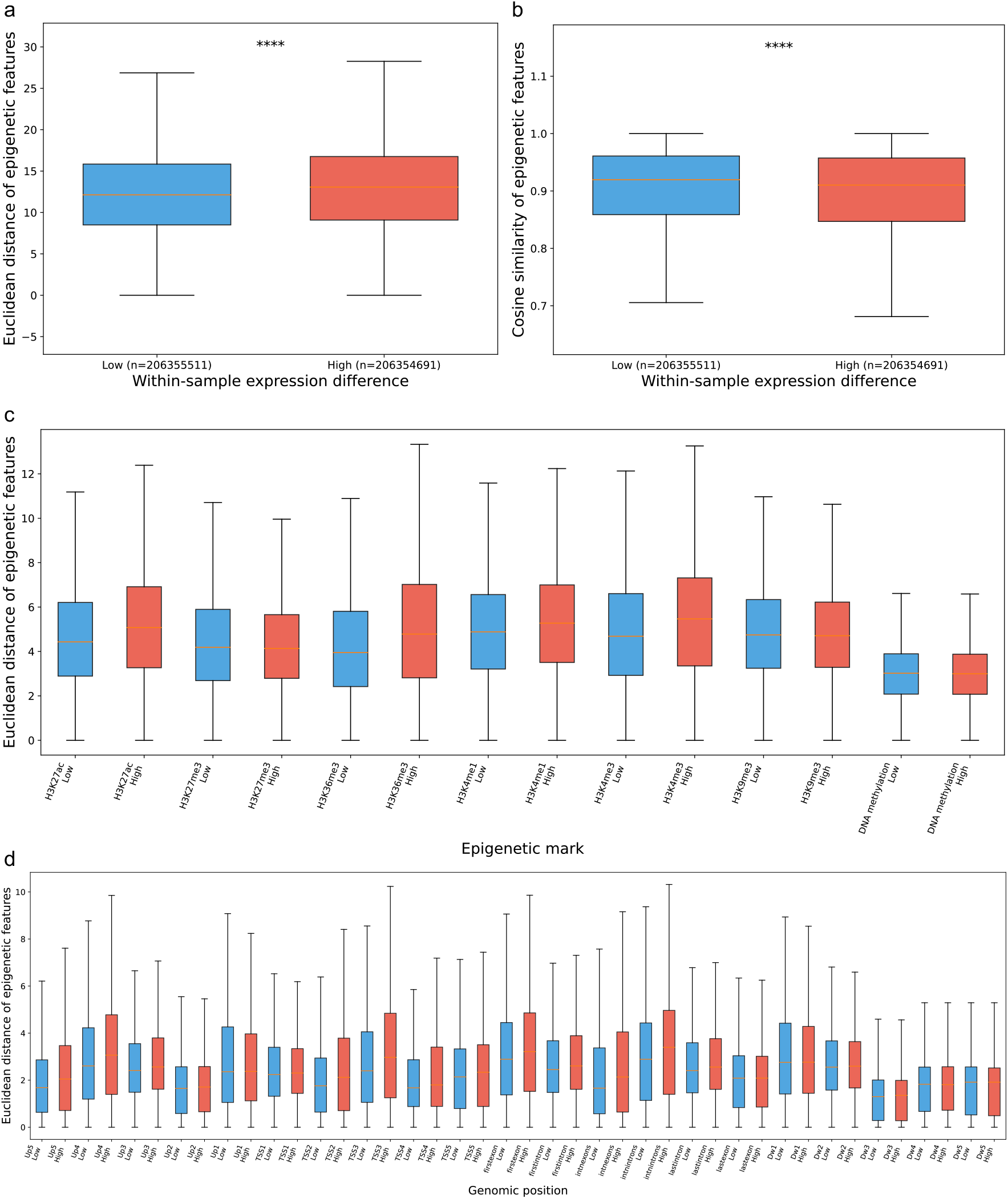
Expression divergence is accompanied by epigenetic divergence between isoforms of the same gene. a, Euclidean distance between paired isoform epigenetic feature vectors stratified by within-sample isoform expression difference. Each observation represents a same-gene isoform pair within an individual biological sample, with low-and high-expression-difference groups defined by the median within-sample expression difference. **b,** Cosine similarity between paired isoform epigenetic feature vectors stratified by within-sample isoform expression difference, reflecting the degree of epigenetic feature redundancy between isoforms. **c,** Mark-specific epigenetic divergence. Euclidean distances were calculated separately for each epigenetic mark across the genomic feature set and compared between low-and high-expression-difference isoform-pair groups. **d,** Position-specific epigenetic divergence. Euclidean distances were calculated separately across genomic feature positions and compared between low-and high-expression-difference isoform-pair groups. Statistical comparisons between groups were performed using two-sided Mann–Whitney U tests. *:p*<*0.05; **:p*<*0.01; ***:p*<*0.001; ****:p*<*0.0001; ns:not significant.

## Supplementary File

Supplementary File S1: List of samples in the core dataset.

## References

[1] Cheng, C. et al. Understanding transcriptional regulation by integrative analysis of transcription factor binding data. Genome Research 22, 1658–1667 (2012).

[2] Tasaki, S., Gaiteri, C., Mostafavi, S. & Wang, Y. Deep learning decodes the principles of differential gene expression. Nature Machine Intelligence 2, 376–386 (2020).

[3] Karbalayghareh, A., Sahin, M. & Leslie, C. S. Chromatin interaction-aware gene regulatory modeling with graph attention networks. Genome Research 32, 930–944 (2022).

[4] Cheng, C. et al. A statistical framework for modeling gene expression using chromatin features and application to modENCODE datasets. Genome Biology 12, R15 (2011).

[5] Dong, X. et al. Modeling gene expression using chromatin features in various cellular contexts. Genome Biology 13, R53 (2012).

[6] Singh, R., Lanchantin, J., Robins, G. & Qi, Y. DeepChrome: Deep-learning for predicting gene expression from histone modifications. Bioinformatics i639–i648 (2016).

[7] Singh, R., Lanchantin, J., Sekhon, A. & Qi, Y. Attend and predict: Understanding gene regulation by selective attention on chromatin. In Advances in Neural Information Processing Systems, vol. 30, 6785–6795 (2017).

[8] Sekhon, A., Singh, R. & Qi, Y. DeepDiff: DEEP-learning for predicting DIFFerential gene expression from histone modifications. Bioinformatics 34, i891—-i900 (2018).

[9] Lee, D., Yang, J. & Kim, S. Learning the histone codes with large genomic windows and three-dimensional chromatin interactions using transformer. Nature Communications 13, 6678 (2022).

[10] Lou, S. et al. Whole-genome bisulfite sequencing of multiple individuals reveals complementary roles of promoter and gene body methylation in transcriptional regulation. Genome Biology 15, 408 (2014).

[11] Li, L., Gao, Y., Wu, Q., Cheng, A. S. L. & Yip, K. Y. New guidelines for DNA methylome studies regarding 5-hydroxymethylcytosine for understanding transcriptional regulation. Genome Research 29, 543–553 (2019).

[12] Seal, D. B., Das, V., Goswami, S. & De, R. K. Estimating gene expression from DNA methylation and copy number variation: A deep learning regression model for multi-omics integration. Genomics 112, 2833–2841 (2020).

[13] Schmidt, F. et al. Combining transcription factor binding affinities with open-chromatin data for accurate gene expression prediction. Nucleic Acids Research 45, 54–66 (2017).

[14] Cao, Q. et al. A unified framework for integrative study of heterogeneous gene regulatory mechanisms. Nature Machine Intelligence 2, 447–456 (2020).

[15] Frasca, F., Matteucci, M., Leone, M., Morelli, M. J. & Masseroli, M. Accurate and highly interpretable prediction of gene expression from histone modifications. BMC Bioinformatics 23, 151 (2022).

[16] Cao, Q. et al. Reconstruction of enhancer-target networks in 935 samples of human primary cells, tissues and cell lines. Nature Genetics 49, 1428–1436 (2017).

[17] Ardakani, F. B., Ashrafiyan, S., Rumpf, L., Hecker, D. & Schulz, M. H. Harnessing machine learning models for epigenome to transcriptome association studies. bioRxiv (2025). URL 10.1101/2025.05.09.653095.

[18] Avsec, Ž., et al. Effective gene expression prediction from sequence by integrating long-range interactions. Nature Methods 18, 1196–1203 (2021).

[19] Avsec, Ž., et al. AlphaGenome: Advancing regulatory variant effect prediction with a unified DNA sequence model. bioRxiv (2025). URL 10.1101/2025.06.25.661532.

[20] Kipf, T. N. & Welling, M. Semi-supervised classification with graph convolutional networks. In Fifth International Conference on Learning Representations (2017).

[21] Bigness, J., Loinaz, X., Patel, S., Larschan, E. & Singh, R. Integrating long-range regulatory interactions to predict gene expression using graph convolutional networks. Journal of Computational Biology 29, 409–424 (2022). URL 10.1089/cmb.2021.0316.

[22] Thurman, R. E. et al. The accessible chromatin landscape of the human genome. Nature 489, 75–82 (2012).

[23] Komarnitsky, P., Cho, E.-J. & Buratowski, S. Different phosphorylated forms of RNA polymerase II and associated mRNA processing factors during transcription. Genes & Development 14, 2452–2460 (2020).

[24] Kelley, D. R. Cross-species regulatory sequence activity prediction. PLoS Computational Biology 16, e1008050 (2020). URL 10.1371/journal.pcbi.1008050.

[25] Linder, J., Srivastava, D., Yuan, H., Agarwal, V. & Kelley, D. R. Predicting rna-seq coverage from dna sequence as a unifying model of gene regulation. Nature Genetics 57, 949–961 (2025). URL 10.1038/s41588-024-02053-6.

[26] Stunnenberg, H. G., Consortium, T. I. H. E. & Hirst, M. The international human epigenome consortium: A blueprint for scientific collaboration and discovery. Cell 167, 1145–1149 (2016).

[27] International Human Epigenome Consortium. EpiATLAS – a reference for human epigenomic research. bioRxiv (2026). URL 10.64898/2026.06.22.729579.

[28] Frankish, A. et al. GENCODE: Reference annotation for the human and mouse genomes in 2023. Nucleic Acids Research 61, DD942–D949 (2023).

[29] Althammer, S., Pagés, A. & Eyras, E. Predictive models of gene regulation from high-throughput epigenomics data. Comparative and Functional Genomics 2012, 284786 (2012). URL 10.1155/2012/284786.

[30] Devailly, G. & Joshi, A. Comprehensive analysis of epigenetic signatures of human transcription control. Molecular Omics 17, 692–705 (2021). URL 10.1039/D0MO00130A.

[31] Merkel, A. et al. gemBS: High throughput processing for DNA methylation data from bisulfite sequencing. Bioinformatics 35, 737–742 (2019).

[32] Oughtred, R. et al. The BioGRID database: A comprehensive biomedical resource of curated protein, genetic, and chemical interactions. Protein Science 30, 187–200 (2020).

[33] Katz, K. et al. The sequence read archive: A decade more of explosive growth. Nucleic Acids Research 50, D387–D390 (2021).

[34] Li, H. & Durbin, R. Fast and accurate short read alignment with burrows–wheeler transform. Bioinformatics 25, 1754–1760 (2009).

[35] Durand, N. C. et al. Juicer provides a one-click system for analyzing loop-resolution hi-c experiments. Cell Systems 3, 95–98 (2016).

[36] Reiff, S. B. et al. The 4D nucleome data portal as a resource for searching and visualizing curated nucleomics data. Nature Communications 13, 2365 (2022).

[37] Kaul, A., Bhattacharyya, S. & Ay, F. Identifying statistically significant chromatin contacts from Hi-C data with FitHiC2. Nature Protocol 15, 991–1012 (2020).

[38] Wang, M., et al. Deep graph library: A graph-centric, highly-performant package for graph neural networks. arXiv (2019). URL https://arxiv.org/abs/1909.01315.

[39] Kingma, D. P. & Ba, J. Adam: A method for stochastic optimization. In Third International Conference on Learning Representations (2015).

[40] Miura, F., Enomoto, Y., Dairiki, R. & Ito, T. Amplification-free whole-genome bisulfite sequencing by post-bisulfite adaptor tagging. Nucleic Acids Reserach 40, e136 (2012).

